# A potent macrocyclic peptide binder enables effector-free crystallization of the GTPase RAB27A

**DOI:** 10.64898/2026.10.13.758952

**Authors:** Rita Petracca, Montse Tersa, Elena De Vita, Thomas Lanyon-Hogg, Yuteng Wu, Anna Kukushkina, Rhodri M. Morgan, Ernesto Cota, Louise J Walport, Edward W Tate

## Abstract

RAB27A is a member of a highly conserved family of small GTPases that regulate intracellular vesicle trafficking. Multiple studies have linked RAB27A cellular activity to exosome-mediated modulation of the tumour microenvironment and inflammatory signalling, highlighting it as an attractive target for drug discovery. However, identifying structurally validated and selective inhibitors of the RAB27A-effector interaction has proven challenging, partly because wild-type RAB27A crystallizes only in complex with effector proteins such as Slp2a. Using mRNA display, we identified macrocyclic peptide ligands that target the RAB27A effector-binding interface. Optimisation of initial hits led to **IMP2660**, a nanomolar-affinity RAB27A binder that effectively disrupts RAB27A-effector interactions. IMP2660 enabled determination of a 2.1 Å structure of active-state RAB27A bound to a noncovalent macrocyclic peptide in the absence of an effector, revealing structural features of the effector-binding surface and providing a platform for future structure-guided ligand development.

## Introduction

RAB27A is a small monomeric G protein (~25 kDa) belonging to the Rab (Ras-like proteins in brain) small GTPase protein family, with about 70 members in humans.^1,2^ Together with Ras, Rho, Ran and Arf, Rab proteins are part of the Ras superfamily, of which they constitute the largest branch. Like other Ras family GTPases, RAB27A cycles between an active, GTP-bound state and an inactive, GDP-bound state (Figure 1).^1,3^ The binding and hydrolysis of GTP induces structural rearrangements in two flexible regions, termed Switch I and Switch II, that form the primary interface for effector recognition. In the active, GTP-bound state, these switch regions are stabilized by the γ-phosphate, creating a rigid surface that is recognized by effector proteins. Conversely, in the GDP-bound form, the Switch regions become disordered and highly dynamic, precluding effector interaction and contributing to the intrinsic instability of isolated Rab proteins.^4^ This inherent structural lability explains why Rab structures are typically determined in complex with effectors and stabilizing GTP analogues.

**Figure 1.**
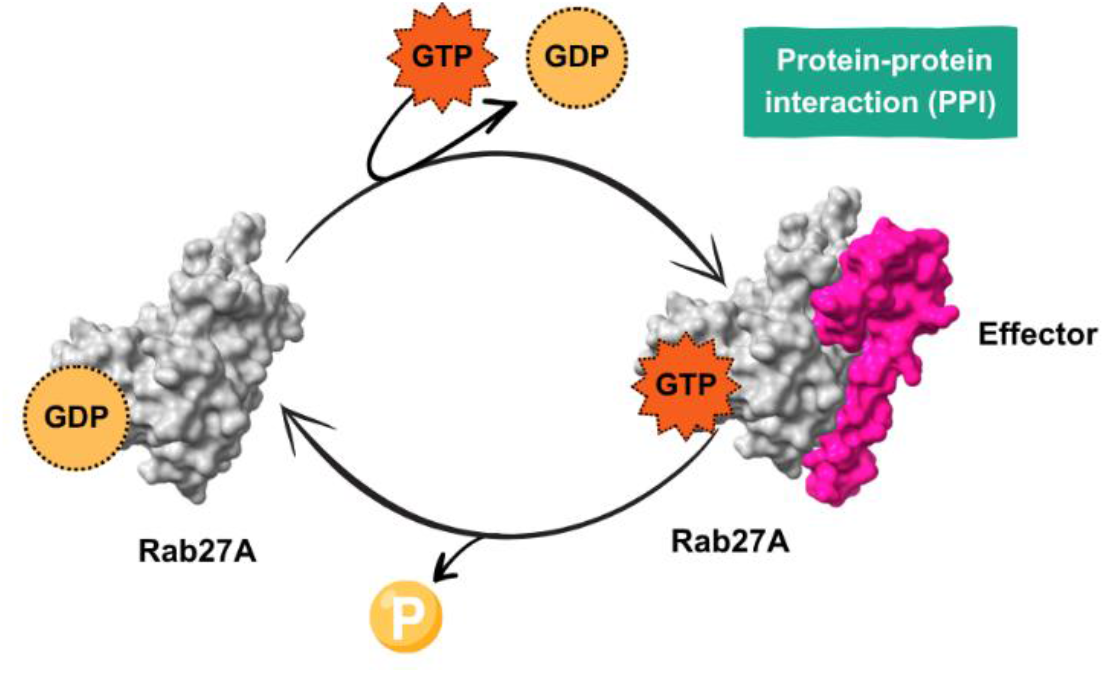
Schematic RAB27A GDP-inactive/GTP active cycle. In the GTP-active state RAB27A (grey) binds the effector proteins, e.g., Slp2a (hot pink, PDB: 3BC1).

Like other Rab GTPases, RAB27 regulates protein transport and membrane trafficking.^5,6^ The two RAB27 isoforms, RAB27A and RAB27B, share >70% similarity in amino acid sequence, and both recruit and interact with various classes of protein effectors, including synaptotagmin-like protein (Slp).^7^ Both isoforms are involved in biogenesis, secretion and distribution of small secreted extracellular vesicles termed exosomes. RAB27A was the first Rab to be associated with a rare heritable condition, Griscelli syndrome type 2 (GS2), which is caused by autosomal recessive loss-of-function mutations.^8^ RAB27-mediated exosome secretion pathways are an emerging cancer hallmark,^9^ and upregulation of RAB27A is associated with malignancy in a range of cancers.^10^

The first Rab27 crystal structure was obtained for mouse (Mus *musculus*) Rab27a in complex with GppNHp nucleotide (GTP analogue), Mg^2+^ ion and a truncated construct of human effector protein Exophilin4 (also termed Slp2A) residues E10 to E62, at 1.8 Å resolution (Figure 2a).^11^ Q78L mouse Rab27a was C-terminally-truncated to residues 1–193 and two additional mutations C123S and C188S were introduced to overcome formation of an intramolecular disulfide bond which impairs co-crystallization with effector.^11^ Rab27a, bound to GTP analogues GppNHp, and Exophilin4 (Slp2A) share an extensive accessible surface area of 1970 Å^2^ with large hydrophobic patches and hydrogen bonding.^11,12^ Rab27a Y122 in the SF4 region interacts via a π-stacking interaction with a W53-Y54 motif in the effector.^11^ This “WF pseudo pocket” is a potential target for inhibition of RAB27-effector protein-protein interactions (PPIs), and alongside the highly conserved nucleotide pocket, it is one of the two predicted ligandable regions in RAB27A. ^13^This pocket is specific to RAB27, and mutation of Y122 strongly inhibits effector binding.^13^

**Figure 2.**
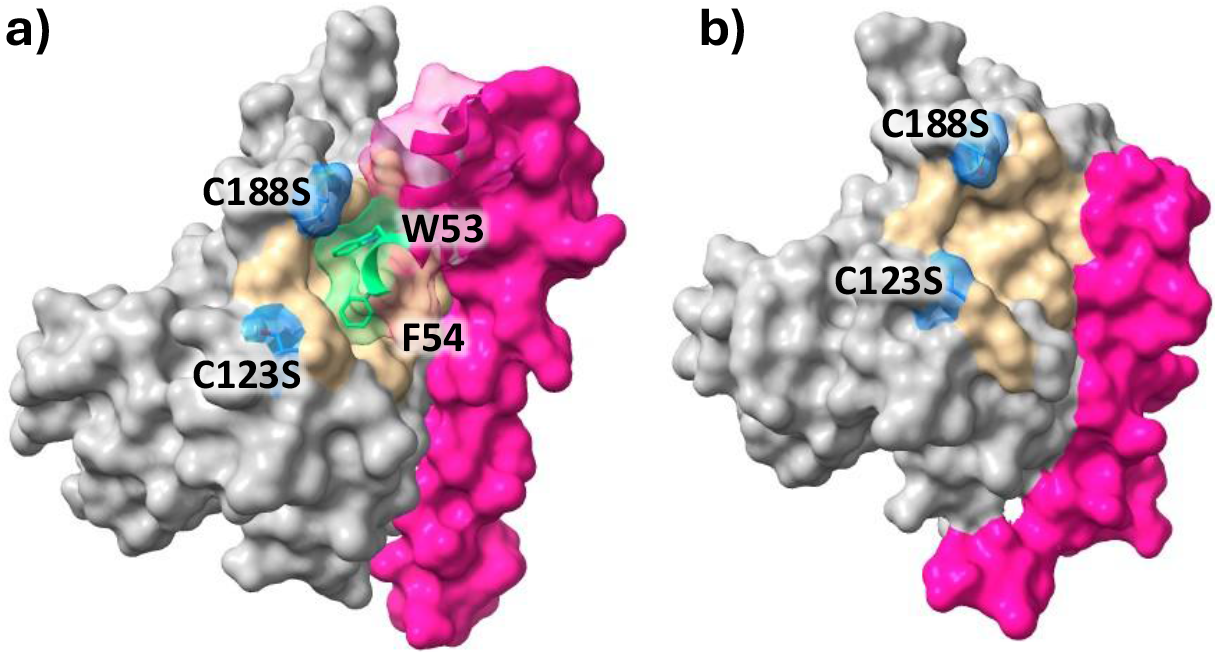
**a)** Previously reported complex of Rab27A(C123S,C188S) (*grey*) with human effector protein SLP2A (residues Glu10 - Glu62), (*pink*) by Chavas *et al*. (PDB: 3BC1),^11^ showing the pseudo-pocket (sand) occupied by W53-Y54 (green); **b)** Our previously reported RAB27A-hSLP2A fusion construct (PDB: 7OPP).^13^

We recently reported the first structurally validated covalent ligands of RAB27A,^13^ identified using the “quantitative Irreversible Tethering” (qIT) assay based on reactivity towards C123 or C188, which flank the WF pseudo-pocket and are unique to RAB27A/B over other Rabs. To allow ligand interactions with the WF site, a portion of effector Slp2A (residues 5–32) was fused to the N-terminus of RAB27A to create a new class of constructs termed RAB-effector fusions (fRAB27A, Figure 2b).^13^ Variants including a C123S and C188S double mutant, as well as mutants named according to retention of cysteine at position 123 (fRAB27A-C123) or 188 (fRAB27A-C188), were expressed and co-crystallized with covalent ligands.^13^

Building on this platform, we recently developed and validated stereoselective Cys123-directed covalent inhibitors of RAB27A/B that engage endogenous RAB27A/B in cells and disrupt RAB27–effector interactions. These findings further establish the WF pocket as a ligandable site and support the development of complementary strategies to pharmacologically modulate the RAB27–effector interface.^14^

Targeting PPI interfaces is intrinsically challenging due to the generally large, flat and hydrophobic surfaces, with no defined pockets or grooves.^15^ Cyclic peptides, due to their high affinity and target specificity have been demonstrated to be suitable for protein-protein interaction (PPI) modulation.^16^ In 2017, a stapled peptide targeting Rab25 was reported.^17,18^

To expand the scope of RAB27-targeted ligand space, we turned our attention towards macrocyclic peptides as potential RAB27A-effector PPI modulators. Herein we report the discovery of the first macrocyclic peptide, **IMP2660**, that efficiently inhibits the RAB27A-Slp2A interaction, as confirmed in a range of biophysical assays. The crystal structure of the RAB27A–**IMP2660** complex provides the first structure of RAB27A bound to a noncovalent macrocyclic peptide in the absence of an effector protein, revealing features of the effector-binding surface that may facilitate future structure-guided ligand development.

## Results and Discussion

### RaPID screening

Macrocyclic peptides can deliver the tight binding characteristic of biologics while targeting challenging PPI interfaces.^19,20^ Large encoded peptide libraries have become powerful tools for discovering high-affinity de novo macrocyclic peptides.^21^ The Random nonstandard Peptides Integrated Discovery (RaPID) system is particularly well suited to this purpose, enabling the generation of structurally diverse libraries containing up to 10^14^ members, including peptides incorporating non-proteinogenic amino acids, and allowing selections under a wide range of experimental conditions. ^22,23,24,25^

In this study, we applied an established RaPID selection workflow^26^ to generate an mRNA-displayed macrocyclic peptide library comprising >10^12^ members. Such library contained six to twelve random amino acids, flanked by an initiator codon and a C-terminal cysteine prior to a GSGSGS-linker. Peptide translation was initiated with *N*-chloroacetyl-D-tyrosine (ClAc-D-Tyr), supplied as its cyanomethyl ester (ClAc-D-Tyr-CME). Compatibility of the activated amino acid with eFx-mediated aminoacylation was assessed using the FAM-labelled microhelix RNA FAM-MiHx_23b, after which ClAc-D-Tyr was charged onto the initiator tRNA for use in the translation system.^27^ Following *in vitro* translation, the *N*-chloroacetyl group underwent spontaneous intramolecular reaction with the C-terminal cysteine to generate the corresponding macrocyclic peptides (§ESI) (Figure 3a).^28^

**Figure 3.**
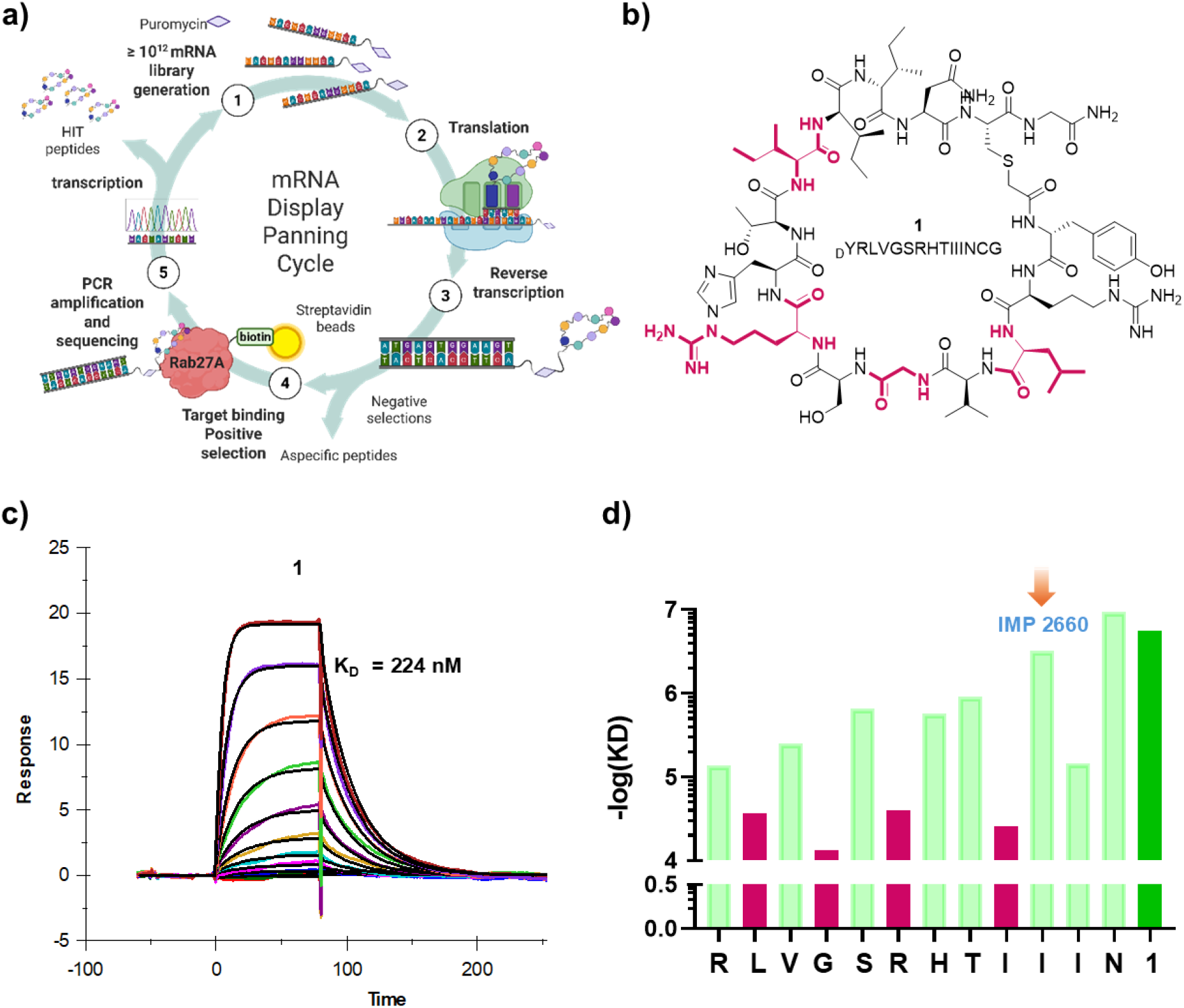
**a)** Schematic depiction of RaPID selection strategy used in this work; **b)** Chemical structure and sequence of peptide **1** and **c)** SPR sensorgram of peptide **1** tested at concentrations of 0, 3.9, 7.5, 15.3, 31.2, 62.5, 125, 250, 500, and 1000 nM. Measurements were performed in 20 mM HEPES, pH 7.4, 150 mM NaCl, 5 mM MgCl_2_, 0.005% Tween-20, and 0.1% BSA. Black lines represent the fits of the experimental sensorgrams to the binding model used for kinetic analysis. **d)** Results of SPR analysis (*n=*3) of **1** single alanine variants at each position as shown on the x axis.

Recombinant RAB27A[C123S/C188S] (residues 1–192, Q78L, GppNHp bound) double mutant, hereafter RAB27A2S, was expressed and biotinylated at the *N*-terminal Gly using Sortase 5M and a biotinylated depsipeptide substrate (§ESI)^13,29^ to allow loading on streptavidin magnetic beads for selection. Five selection cycles led to considerable enrichment of sequences in the initial library, when analyzed by next generation sequencing, and identified several potential hits which were considered for further preliminary analysis (§ESI).

### Synthesis and biological evaluation of selected macrocyclic peptides

Selected peptides were synthesized according to standard SPPS protocols (§ESI) and evaluated for binding to biotinylated RAB27A2S by Surface Plasmon Resonance (SPR) using a streptavidin-coated chip. During the initial screening, several peptides showed substantial bulk refractive index effects, whereas peptide **1** produced a cleaner concentration-dependent binding response and was therefore selected for further characterization. Under optimized SPR conditions, peptide **1** exhibited a K^D^ of 224 nM and was taken forward as the starting point for subsequent investigations (Figure 3b).

As a first step to define the key determinants of binding, we performed an alanine-scanning analysis^29,30^ across the peptide sequence and quantified binding affinities by SPR (§ESI) (Figure 3d) Substitution of individual residues with alanine revealed distinct contributions to target recognition. In general, substitutions in the *N*-terminal and central motifs caused a pronounced loss of affinity (Δlog K^D^ > 1), whereas C-terminal variants were more tolerated. Replacement of **1**L2, **1**G4, **1**R6, and **1**I9 residues markedly reduced binding, suggesting an important role in binding. Interestingly, IMP2660 bearing an A substitution at position 10 displayed markedly improved solubility and chromatographic behavior relative to the parent peptide, potentially due to disruption of a hydrophobic patch formed by the Ile triplet at positions 9–11. Since IMP2660 retained high target affinity and could be obtained in greater yield and purity (§ESI), this analogue was selected for further functional studies. Binding of IMP2660 to the free protein in solution was further confirmed using an orthogonal differential scanning fluorimetry (DSF)^31,32^ assay with RAB27A2S (Figure 4a). At the highest tested concentration, IMP2660 induced a 4.91 °C increase in the melting temperature (ΔTm) of RAB27A2S, consistent with protein stabilization upon ligand binding.

**Figure 4.**
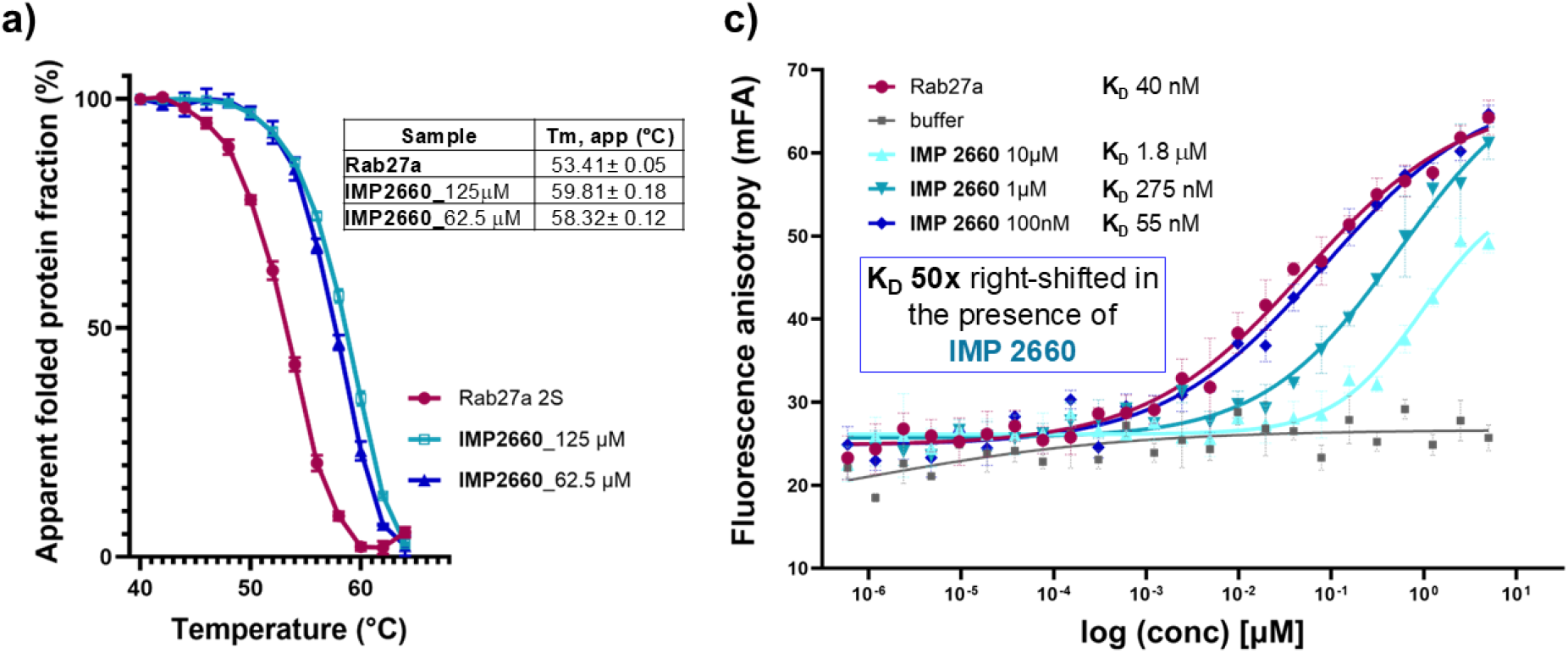
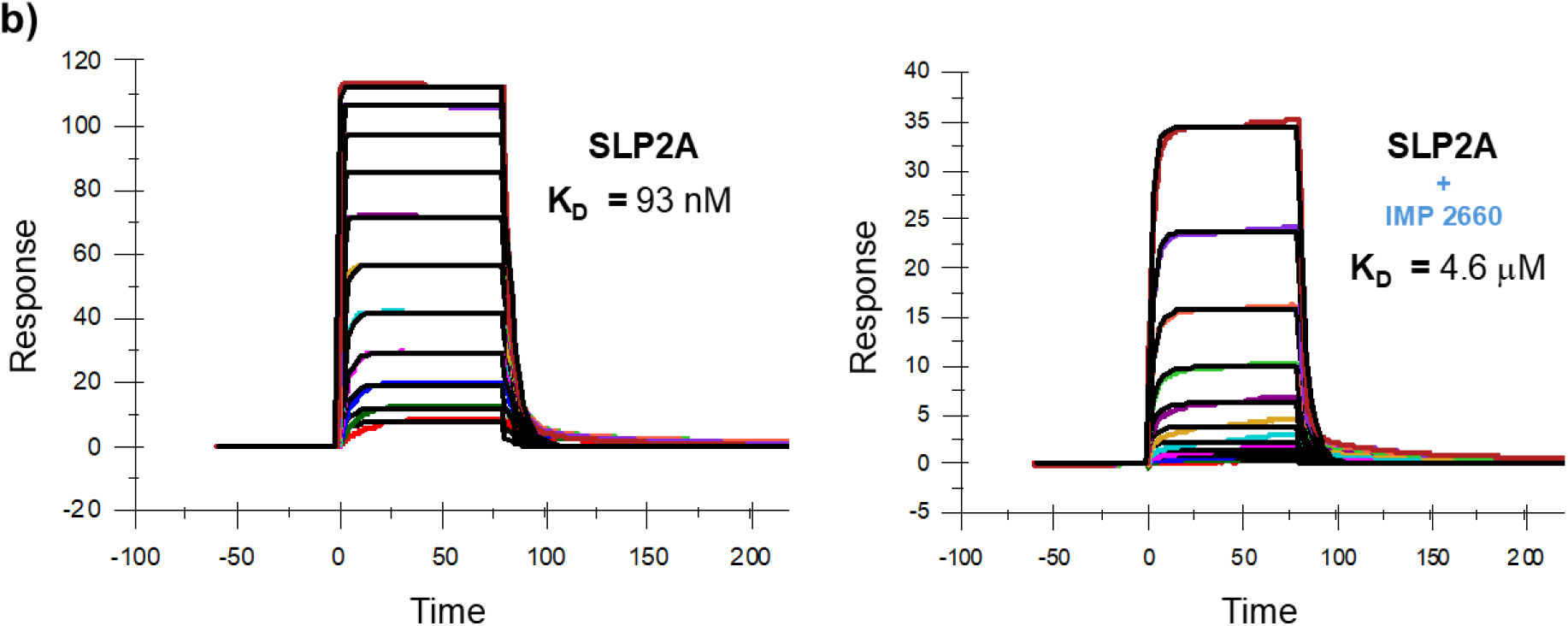
**a)** DSF assay showing a concentration-dependent thermal stabilization of RAB27A2S upon IMP2660 binding (ΔT_m_ = 4.91 °C at 62.5 µM); **b)** SPR sensorgrams of SLP2A binding RAB27A in the presence (K_D_ = 4.6 µM) or absence (K_D_ = 93 nM) of 10 µM IMP2660; **c)** FP competition assay showing a 50-fold right-shift in K_D_ in the presence of IMP2660, consistent with direct competition for the SLP2A-binding site.

### IMP2660 disrupts PPI with effector protein Slp2A

Having confirmed IMP2660 as a potent RAB27A2S binder, we next focused on evaluating its potential as an effective PPI inhibitor. Recombinant truncated SLP2A (E10–E62) (§ESI) was expressed and purified and used to mimic the full effector protein for assessment of binding affinity. SPR analysis confirmed that SLP2A bound to RAB27A2S with a K^D^ of 93 nM (Figure 4b). Under the same conditions, IMP2660 showed a comparable binding affinity of 96 nM (§ESI). To determine whether IMP2660 was capable of displacing SLP2A as a potential PPI inhibitor we established a competitive SPR assay (§ESI). *N*-terminal biotinylated RAB27A2S was immobilized on a streptavidin chip, and the binding affinity of SLP2A was tested in a buffer containing a fixed concentration of IMP2660 (§ESI). The second sensorgram reported in Figure 4b shows a marked decrease in SLP2A affinity in the presence of the peptide, consistent with competitive occupancy of the SLP2A-binding site by IMP2660. Competition was further confirmed in solution using a fluorescence polarization (FP) assay. SLP2A was *N*-terminally labeled with a TAMRA fluorophore (TAMRA–SLP2A) using Sortase 5M and a TAMRA-containing depsipeptide (§ESI). RAB27A2S was titrated in the presence of a fixed concentration of TAMRA– SLP2A, previously optimized for this assay, with binding monitored through an increase in FP signal (§ESI) (Figure 4c). Upon increasing concentrations of IMP2660, a pronounced right-shift in the binding curve was observed, corresponding to an approximately 50-fold reduction in effector binding affinity at 10 µM IMP2660. Together, these data confirm that IMP2660 binds specifically and with high affinity to RAB27A in competition with the effector, significantly stabilizing the protein.

### IMP2660 permits crystallization of RAB27A2S in the absence of effector

We next sought to confirm the binding mode of IMP2660 by X-ray crystallography. Crystallization of minimally engineered RAB27A in the absence of an effector remains challenging, likely due to the conformational flexibility of the unbound protein and the lack of intermolecular contacts required for efficient crystal packing. We previously reported a structure of RAB27A bearing multiple mutations to engineer crystal contacts, but this approach brought disadvantages including highly complex crystal symmetry.^30^ Given these significant challenges, we were delighted to discover that in the presence of **IMP2660**, RAB27A2S could be crystallized and the complex solved by X-ray crystallography to 2.1 Å with minimal optimization of previously-reported conditions (§ESI). The asymmetric unit of this structure includes two Rab27A2S chains (A and B, Figure 5a) and closely resembles the dimers observed in our previously-reported RAB27A-SLP2a fusion construct (PDB: 7OPP)^13^ and the effector-bound complex (PDB: 3BC1),^11^ which is unsurprising given that recombinant RAB27A2S bears a Q78L mutation and is GppNHp bound, ensuring a GTP-bound conformation. This also implies that IMP2660 acts as a surrogate for truncated effector and is able to confer sufficient stability to RAB27A2S to achieve crystallization of the active conformation. The electron density map confirms the presence of one molecule of IMP2660 bound to each chain, with defined density for the macrocycle portion of the molecule which engages in stable contacts with Rab27A2S, and unresolved flexibility observed only for the *N*-terminal glycine.

**Figure 5.**
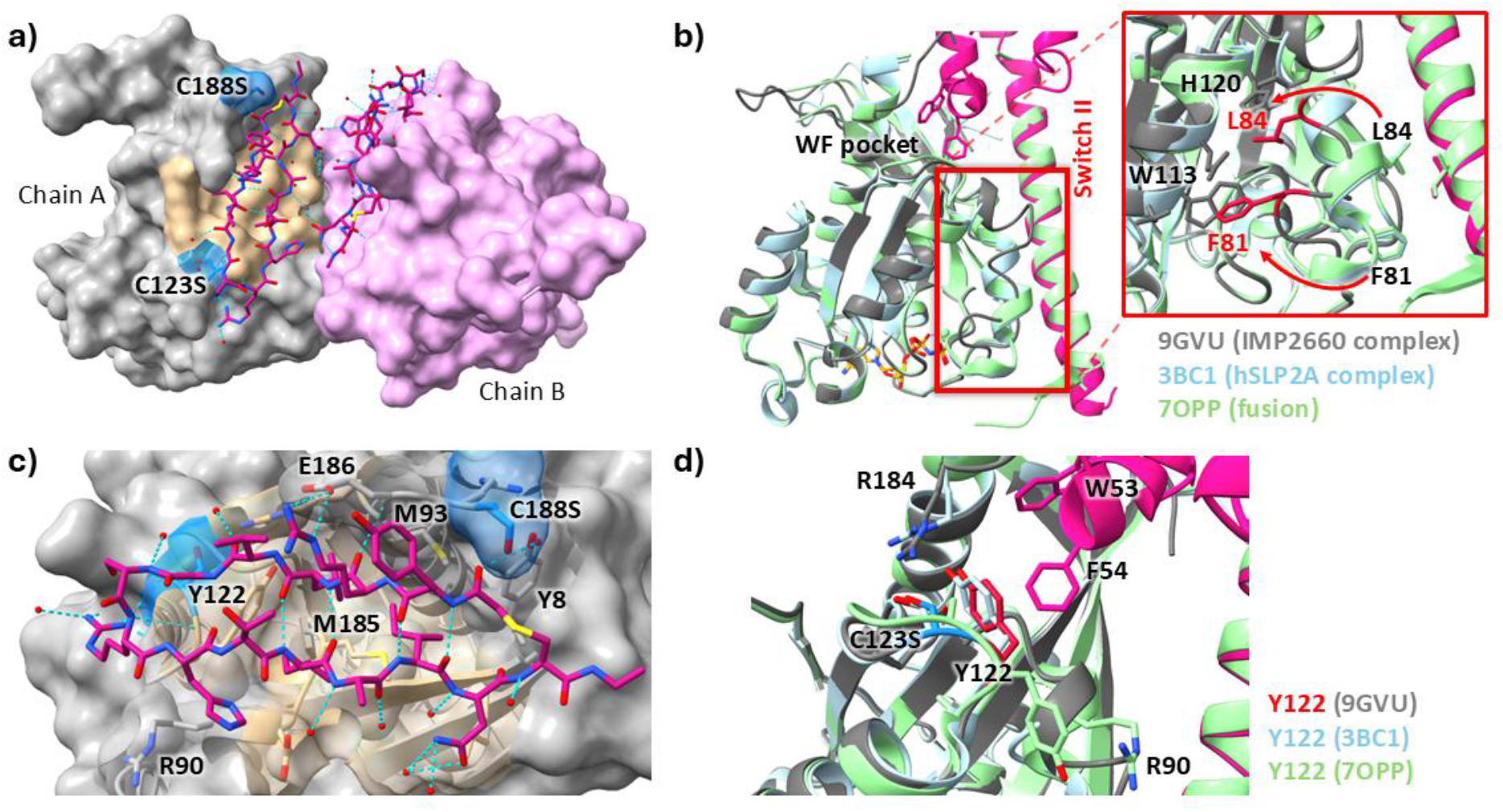
**a)** RAB27A2S dimer in complex with **IMP2660** (*magenta*, PDB: 9GVU), showing both chain A (*grey*) and B (*lilac*); **b)** Superimposition of the effector-free structure (9GVU, *grey*) to RAB27A (*cyan*) in complex with SLP2A (*pink*, PDB: 3BC1) and our previously reported SLP2A-RAB27A fusion (PDB: 7OPP). Red square highlights the differences in the Switch II region and zoomed detail on the right panel shows F81 and L84 (*red*) introversion compared to SLP2A-containing structures; **c)** Close-up view of the main interactions between **IMP2660** (*magenta*) and RAB27A2S (Chain A, *grey*); **d)** Conformation of Y122 in the **IMP2660**-RAB27A2S complex (highlighted in *red*) resembles the inward orientation of the observed when in complex with SLP2A (*cyan*). In the fusion construct (*green*), lacking the WF motif, Y122 reorients towards Arg90 of the WF pocket.

The spatial organization of the Rab27A2S dimer is not identical to earlier structures, but the single chains have good superimposition with our previous model (PDB: 7OPP), with an average RMSD of 1.8 Å. However, all interactions between RAB27A2S and IMP2660 occur in the WF pocket, and the α-helicity of the switch II region (the short helix α2, L78-A92), which is observed as a partial loop in this structure, is reduced in comparison to the structured helix observed with SLP2A (Figure 5b). Major rearrangements in this part of the structure are driven by introversion of F81 and L84 to escape solvent contacts. This rearrangement causes loss of α-helix conformation compared to effector-bound structures and may represent the conformational state of GTP-bound RAB27A prior to effector binding.

The two RAB27A2S molecules (chain A and B) have an RMSD of 1.0 Å, and the two bound **IMP2660** molecules have a RMSD of 1.1 Å, showing an overall consistent binding mode within the asymmetric unit. While the dimer interface includes the WF binding site, contacts between the two macrocyclic molecules are minimal and mostly bridged via water, implying that the observed binding pose is driven by affinity for RAB27A2S. The most important interactions engaged within both chain A and B include hydrophobic contacts with M93 and M185 by the V3 and both I9 and I11 residues of **IMP2660** (Figure 5c). Interestingly, these contacts closely recapitulate the binding of the WF residues from effectors at this site, with Y122 remaining in an inward conformation, as opposed to the outward conformation observed with fusion constructs lacking the WF binding motif (Figure 5d). However, no π-π interaction to Y122 is engaged by **IMP2660** which occupies this site with the I9 residue^.13^

The hydrophilic residues of **IMP2660** engage in a network of hydrogen bonding mainly driven by the two R1 and R6 residues that have multiple interactions each with the acidic residue patch 118–123 on RAB27A2S. In particular, the lateral chain of E186 significantly moves to point toward the WF pocket, where it can engage the R1 of IMP2660 via H-bonding (as seen in chain A, Figure 5a). The H7 residue forms H-bonds with the backbone amide of R90, and the N12 with L9, consolidating the conformation of the macrocycle which is supported also by the intramolecular contacts of the peptidic backbone. Overall, this well-defined binding pose is consistent with the affinity measured in biophysical assays and the results from alanine-scanning and provides evidence for direct competition with effector interaction at the WF pocket.

## Conclusion

In this work we investigated macrocyclic peptides as potential inhibitors of the protein–protein interaction (PPI) between RAB27A and its effector proteins, demonstrating the feasibility and success of this approach for RAB27A. A potent peptide binder (**IMP2660**) was identified which binds RAB27A2S with a K^D^ of 96 nM (by SPR) and it specifically targets the RAB27A–SLP2A interaction at the SF4 domain. Potency was assessed by orthogonal biophysical and biochemical assays, including a novel fluorescence polarization assay using fluorescently labelled SLP2A. Binding kinetics and effector displacement were verified by SPR. The specific binding mode of **IMP2660** was further elucidated by X-ray crystallography, enabling determination of the structure of active-state RAB27A bound to a noncovalent macrocyclic peptide in the absence of an effector protein, with minimal mutations. The structural elucidation of the active form of RAB27A bound to **IMP2660** revealed a more flexible switch II region (the short helix α2, L78-A92) compared to previously reported effector complexes, which hints at this region being critical for specific effector recognition. This is in line with the importance of this region observed for the function of other Ras superfamily GTPases, and may imply specific effector-binding dynamics for the active form of RAB27 GTPases.^31^ Furthermore, compared with covalent small molecules, IMP2660 engages Y122 in an inward orientation, recapitulating the binding conformation observed with SLP2A.^13^ IMP2660 therefore represents a complementary tool to small-molecule RAB27A inhibitors. Beyond its use as a structural ligand, a fluorescently labelled IMP2660 analogue has recently been employed as a high-affinity FP probe for interrogating ligand binding at the RAB27A effector interface.^14^ Together, these findings establish IMP2660 as a structurally validated chemical tool for probing RAB27A–effector interactions and provide a starting point for the development of next-generation ligands suitable for interrogating RAB27A-mediated pathways in cellular systems.

## Supporting Information

The authors have cited additional references within the Supporting Information.

## Acknowledgements

We thank Diamond Light Source for access to beamline I02 (proposal No. mx17221 and mx23620), which contributed to the results presented here. The X-ray facility at Imperial College was funded by BBSRC (BB/D524840/1) and the Wellcome Trust (202926/Z/16/Z). The authors would like to thank L. Haigh for managing the Imperial College Chemistry Department Mass Spectrometry suite and P. Haycock for managing the Imperial College Chemistry Department NMR facility. RP is supported by a H2020 (EC) MSCA-IF, project ProCenDecl (EU project 797995). EDV is also supported by a H2020 (EC) MSCA-IF, project RabTarget4Metastasis (EU project 890900). This work was supported by the Francis Crick Institute, which receives its core funding from Cancer Research UK (CC2030), the UK Medical Research Council (CC2030) and the Wellcome Trust (CC2030)

