## Supplementary Information for "A potent macrocyclic peptide binder enables effector-free crystallization of the GTPase RAB27A"

**for**

### Table of contents


| | |
|---|---|
| **Supplementary Figures** | **S3** |
| **Supplementary Tables** | **S6** |
| **Abbreviations** | **S9** |
| **Materials and Methods** | **S10** |
| **Appendix** | **S15** |
| **Supplementary References** | **SX** |

### Supplementary Figures

**Figure S1.** Recovery and enrichment of the RAB27A2S-binding RaPID library across the five selection rounds. Plot recovered library relative to input and the negative-selection control for each round.

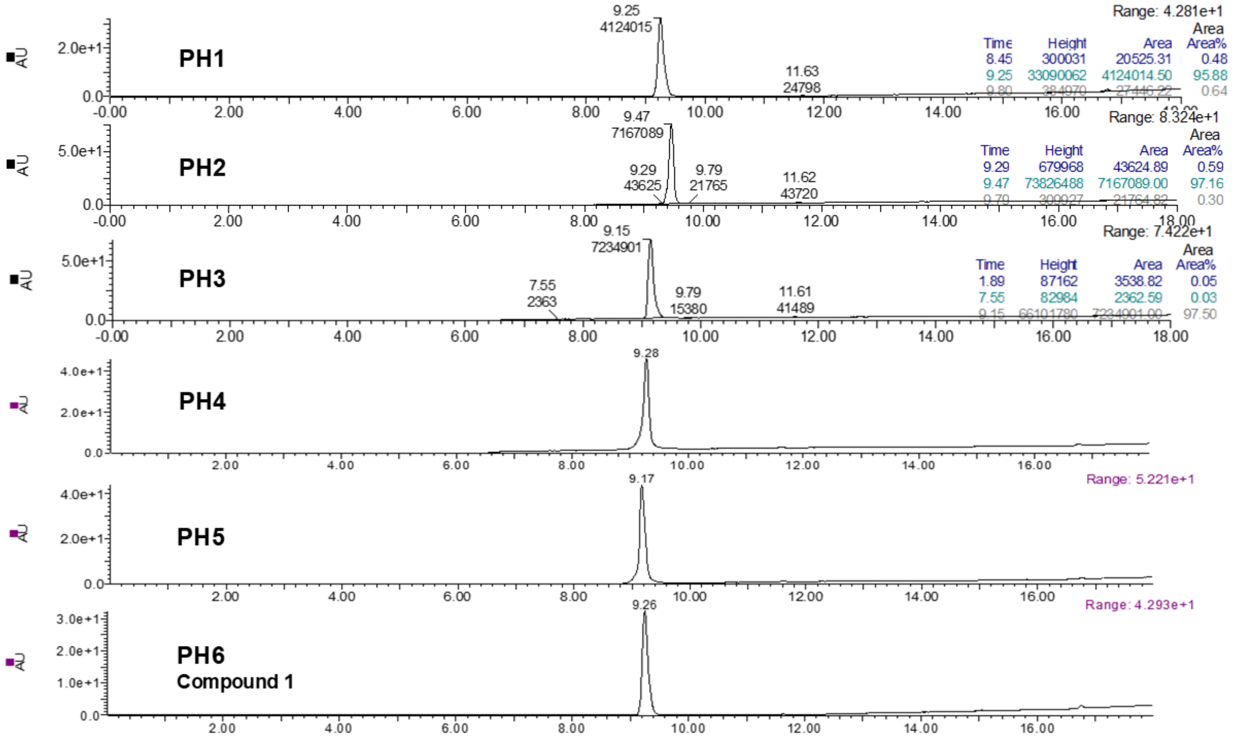


**Figure S2.** LC-MS traces of HPLC purified cyclic peptide hits (PH1-PH6). The calculated and observed mass values are reported in Table S1.

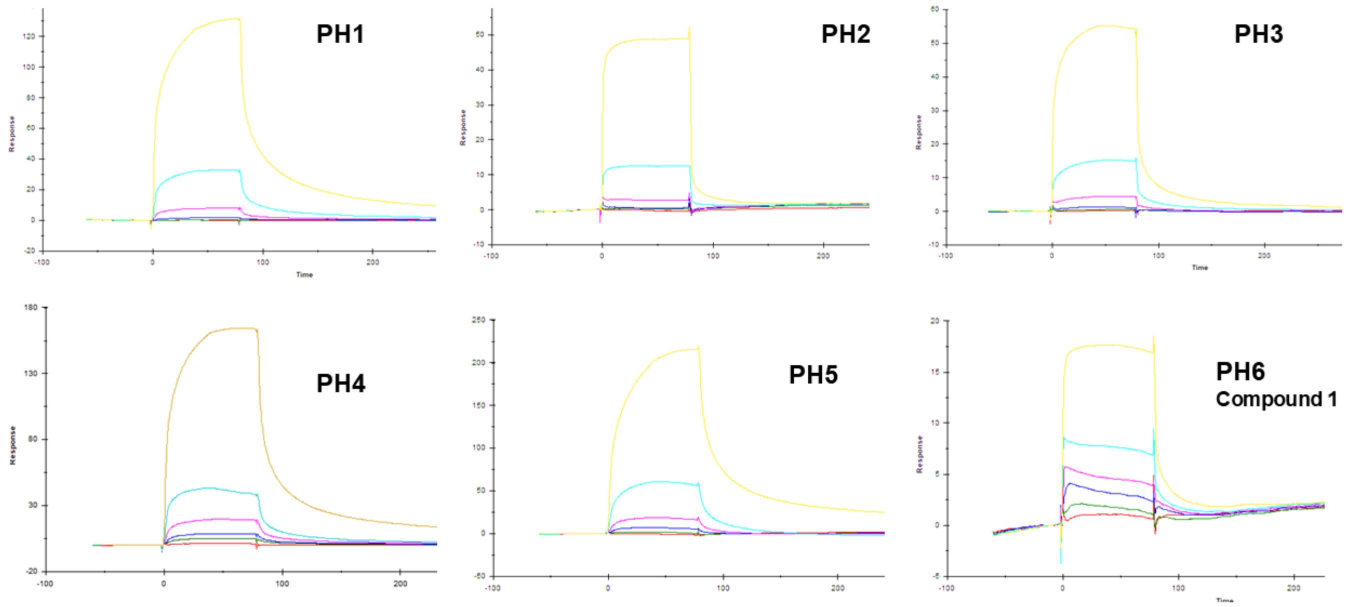


**Figure S3.** Representative SPR sensorgrams of purified cyclic peptide hits PH1–PH6 binding to immobilized biotinylated RAB27A2S. Peptides were tested at concentrations of 0, 62.5, 125, 250, 500, 1000, and 4000 nM. Measurements were performed in 20 mM HEPES, pH 8.0, 100 mM NaCl, 5 mM $MgCl_2$, and 0.005% Tween-20. PH1–PH5 showed a pronounced bulk refractive index effect, whereas PH6 (compound 1) did not display the same effect.

**Figure S4.** Representative SPR sensorgrams for the single-alanine variants of peptide 1.

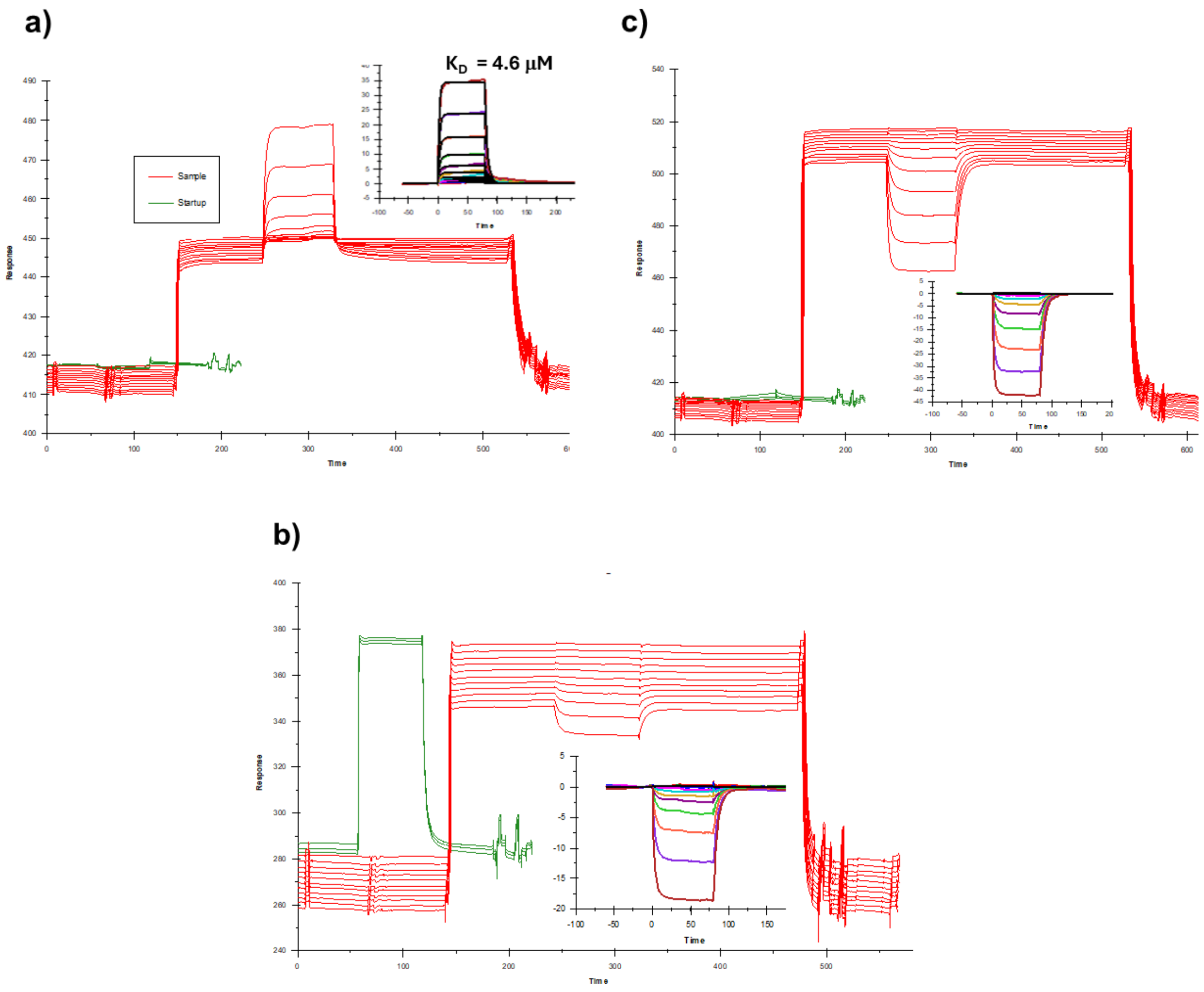


**Figure S5.** SPR competition analysis of IMP2660 and SLP2A binding to bio-RAB27A2S. Competition experiments were performed using surface-immobilised biotinylated RAB27A and an A–B–A injection format. The flanking injections contained a) 10 µM IMP2660, b) 10 µM SLP2A or c) 50 µM SLP2A, with increasing concentrations of the competing ligand. Full raw SPR sensorgrams and the corresponding processed competition responses (insets) are shown. In the presence of 5 µM IMP2660, the apparent dissociation constant for SLP2A was 4.6 µM. Competition and individual binding experiments were performed in running buffer containing 20 mM HEPES (pH 7.4), 150 mM NaCl, 5 mM $MgCl_2$, 0.005% Tween 20 and 0.1% BSA. Responses are expressed in resonance units (RU).

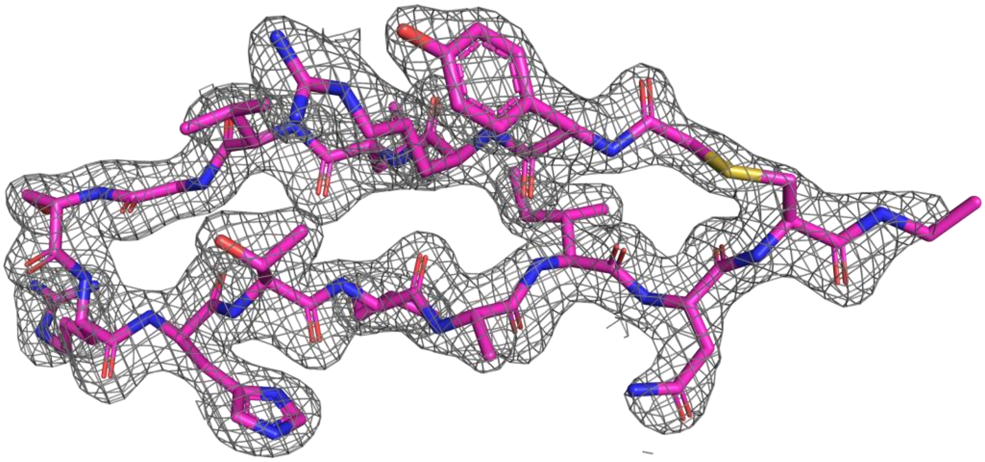

**Figure S6.** Electron-density maps for **IMP2660** bound to chains A and B of RAB27A2S (PDB 9GVU). $2F_o$-Fc maps contoured at 1 σ around the macrocycle.

#### Supplementary Tables

**Table S1.** Peptide hits (PH) obtained after 5 rounds of RaPID selection. The cyclic peptides were synthesized according to standard SPPS protocols, purified by HPLC and analyzed by LC-MS. The LC-MS traces are reported in Supplementary Figure S2.

| Peptide | Linear precursor sequence | Chemical Formula | $[M+H]^+$ | $[M+2H]^{2+}$ | $[M+3H]^{3+}$ | Observed mass |
|---|---|---|---|---|---|---|
| **PH1** | ClAc-DY-CLIDFKHLFCR-$NH_2$ | $C_{75}H_{109}N_{19}O_{16}S_2$ | 1596.78 | 798.89 | 532.93 | 1597/799 |
| **PH2** | ClAc-DY-WLIDFKHLFCR-$NH_2$ | $C_{83}H_{114}N_{20}O_{16}S$ | 1679.85 | 840.43 | 560.62 | 1680/841 |
| **PH3** | ClAc-DY-LIDFKHLFCR-$NH_2$ | $C_{81}H_{113}N_{19}O_{17}S$ | 1656.83 | 828.92 | 552.95 | 1657/829 |
| **PH4** | ClAc-DY-LIDFKHLFCRCG-$NH_2$ | $C_{86}H_{121}N_{21}O_{19}S_2$ | 1816.86 | 908.93 | 606.29 | 1817/909 |
| **PH5** | ClAc-DY-LIDFKHLFCRCG-$NH_2$ | $C_{86}H_{121}N_{21}O_{19}S_2$ | 1816.86 | 908.93 | 606.29 | 1817/909 |
| **PH6 (1)** | ClAc-DY-RLVGSRHTIIINCG-$NH_2$ | $C_{76}H_{125}N_{25}O_{20}S$ | 1740.93 | 870.96 | 580.98 | 871.66/581.56 |

**Table S2.** Sequences and RAB27A2S-binding affinities of the parent peptide and optimised macrocycle. Cyclisation occurs by thioether formation between the *N*-terminal chloroacetyl group and the cysteine side chain. *Of cyclic peptide after HPLC purification.

| Peptide | Linear precursor sequence | Substitution | Yield (%)* | KD by SPR |
|---|---|---|---|---|
| **1** | ClAc-DY-RLVGSRHTIIINCG-$NH_2$ | Parent | 5 | 224 nM |
| **IMP2660** | ClAc-DY-RLVGSRHTIAINCG-$NH_2$ | I10A | 15 | 96 nM |

**Table S3.** Single-alanine variants of peptide 1. Yields of the corresponding cyclic peptides after HPLC purification and KD values obtained by SPR evaluation (as a means of three independent measurements).

| Variant | Linear precursor sequence | Chemical formula | $[M+H]^+$ | $[M+2H]^{2+}$ | $[M+3H]^{3+}$ | Observed mass | $K_D$ by SPR |
|---|---|---|---|---|---|---|---|
| **R1A** | ClAc-DY-**A**LVGSRHTIIINCG-$NH_2$ | $C_{73}H_{118}N_{22}O_{20}S$ | 1655.86 | 828.44 | 552.63 | | 7.26 µM |
| **L2A** | ClAc-DY-R**A**VGSRHTIIINCG-$NH_2$ | $C_{73}H_{119}N_{25}O_{20}S$ | 1698.88 | 849.95 | 566.97 | | 26.9 µM |
| **V3A** | ClAc-DY-RL**A**GSRHTIIINCG-$NH_2$ | $C_{74}H_{121}N_{25}O_{20}S$ | 1712.90 | 856.95 | 571.64 | 857.53/572.13 | 3.98 µM |
| **G4A** | ClAc-DY-RLV**A**SRHTIIINCG-$NH_2$ | $C_{77}H_{127}N_{25}O_{20}S$ | 1754.94 | 877.98 | 585.65 | 878.59/585.81 | 74.7 µM |
| **S5A** | ClAc-DY-RLVG**A**RHTIIINCG-$NH_2$ | $C_{76}H_{125}N_{25}O_{19}S$ | 1724.94 | 862.97 | 575.65 | 863.69/575.78 | 1.51 µM |
| **R6A** | ClAc-DY-RLVGS**A**HTIIINCG-$NH_2$ | $C_{73}H_{118}N_{22}O_{20}S$ | 1655.86 | 829.94 | 553.63 | | 25 µM |
| **H7A** | ClAc-DY-RLVGSR**A**TIIINCG-$NH_2$ | $C_{73}H_{123}N_{23}O_{20}S$ | 1674.91 | 837.95 | 559.97 | | 1.75 µM |
| **T8A** | ClAc-DY-RLVGSRH**A**IIINCG-$NH_2$ | $C_{75}H_{123}N_{25}O_{19}S$ | 1710.92 | 855.96 | 570.98 | 856.01/570.99 | 1.10 µM |
| **I9A** | ClAc-DY-RLVGSRHT**A**IINCG-$NH_2$ | $C_{73}H_{119}N_{25}O_{20}S$ | 1698.88 | 849.94 | 566.97 | 849.86/566.74 | 39.7 µM |
| **I10A** | ClAc-DY-RLVGSRHTI**A**INCG-$NH_2$ | $C_{73}H_{119}N_{25}O_{20}S$ | 1698.88 | 849.94 | 566.97 | 850.68/567.57 | 0.313 µM |
| **I11A** | ClAc-DY-RLVGSRHTII**A**NCG-$NH_2$ | $C_{73}H_{119}N_{25}O_{20}S$ | 1698.88 | 849.94 | 566.97 | | 6.8 µM |
| **N12A** | ClAc-DY-RLVGSRHTIII**A**CG-$NH_2$ | $C_{75}H_{124}N_{24}O_{19}S$ | 1697.93 | 849.47 | 566.65 | 1699 | 0.105 µM |

**Table S4.** Data-collection and refinement statistics for human RAB27A2S in complex with IMP2660 (PDB 9GVU). Statistics for the highest-resolution shell are shown in parentheses.

| Parameter | Value |
|---|---|
| **Data collection** | |
| Space group | P 21 21 21 |
| Unit-cell parameters (Å) | a = 70.357, b = 72.714, c = 91.658; α = β = γ = 90° |
| Wavelength (Å) | 0.9999 |
| Resolution range (Å) | 56.97-2.10 (2.175-2.10) |
| Total reflections | 352,887 (32,648) |
| Unique reflections | 28,093 (2,759) |
| Multiplicity | 12.5 (12.0) |
| Completeness (%) | 99.80 (98.81) |
| Mean I/σ(I) | 4.39 (0.56) |
| Rmerge | 0.2523 (1.121) |
| Rmeas | 0.2636 (1.17) |
| Wilson B factor ($Å^2$) | 41.8 |
| CC1/2 | 0.951 (0.821) |
| **Refinement** | |
| Reflections used in refinement | 28,049 (2,730) |
| Reflections used for Rfree | 1,398 (114) |
| Rwork | 0.1887 (0.2716) |
| Rfree | 0.2308 (0.3202) |
| RMSD bond lengths (Å) | 0.008 |
| RMSD bond angles (°) | 0.92 |
| **Average B factor ($Å^2$) / atom count** | |
| Macromolecules | 44.93 / 2,830 |
| Water molecules | 52.17 / 175 |
| Ligands and ions | 45.89 / 383 |
| Ramachandran favoured (%) | 95.84 |
| Ramachandran allowed (%) | 3.88 |
| Ramachandran outliers (%) | 0.28 |
| Rotamer outliers (%) | 0.35 |

**Table S4.** Apparent melting temperatures of RAB27A2S in the presence and absence of IMP2660.

| Sample | Apparent Tm (°C) | ΔTm (°C) |
|---|---|---|
| RAB27A2S | 53.41 ± 0.05 | - |
| RAB27A2S + 62.5 μM IMP2660 | 58.32 ± 0.12 | +4.91 |
| RAB27A2S + 125 μM IMP2660 | 59.81 ± 0.18 | +6.40 |

**Table S5.** Apparent affinity of RAB27A2S for TAMRA-SLP2A measured by fluorescence polarisation at fixed IMP2660 concentrations. The values of KD are reported as means of three different measurements.

| IMP2660 concentration | Apparent KD | Fold shift versus control |
|---|---|---|
| 0 | 40 nM | 1.0 |
| 100 nM | 55 nM | 1.4 |
| 1 μM | 275 nM | 6.9 |
| 10 μM | 1.8 μM | 45 (~50) |

**Table S6.** Protein constructs used in this study.

| Construct | Sequence boundaries | Mutations or tags | Use |
|---|---|---|---|
| RAB27A2S | Human RAB27A 1-192 | Q78L, C123S, C188S; N-terminal $His_6$-TEV | RaPID, SPR, DSF, FP, crystallography |
| SLP2A | Human SLP2A 5-62 | N-terminal $His_6$-GB1-TEV | SPR and TAMRA-FP tracer |

#### Abbreviations

**BSA:** bovine serum albumin
**ClAc:** *N*-chloroacetyl
**CME:** cyanomethyl ester
**DCM:** dichloromethane
**DIPEA:** *N*,*N*-diisopropylethylamine
**DMF:** *N*,*N*-dimethylformamide
**DMSO:** dimethyl sulfoxide
**DSF:** differential scanning fluorimetry
**DTT:** dithiothreitol
**ESI:** electronic supplementary information
**FIT:** flexible *in vitro* translation
**FP:** fluorescence polarisation
**GppNHp:** guanosine 5′-(β,γ-imido)triphosphate
**HPLC:** high-performance liquid chromatography
**IPTG:** isopropyl β-D-1-thiogalactopyranoside
**LC-MS:** liquid chromatography-mass spectrometry
**MES:** 2-(*N*-morpholino)ethanesulfonic acid
**NGS:** next-generation sequencing
**NTA:** nitrilotriacetic acid
**PPI:** protein-protein interaction
**RaPID:** Random nonstandard Peptides Integrated Discovery
**RU:** response unit
**SDS-PAGE:** sodium dodecyl sulfate-polyacrylamide gel electrophoresis
**SPPS:** solid-phase peptide synthesis
**SPR:** surface plasmon resonance
**TAMRA:** carboxytetramethylrhodamine
**TEV:** Tobacco Etch Virus
**TFA:** trifluoroacetic acid
**TIPS:** triisopropylsilane
**Tris:** tris(hydroxymethyl)aminomethane

### Materials and Methods

#### ▪ General information

Unless stated otherwise, reagents were obtained from commercial suppliers including Merck, Cytiva Life Science, New England Biolabs (NEB) and Invitrogen and used without further purification. Fmoc-D-Tyr(tBu)-OH was obtained from Fluorochem (Hadfield, UK). Fmoc-Cys(4-methoxytrityl)-OH (Cys-Mmt) was obtained from Sigma Aldrich. All other Fmoc protected amino acids were obtained from AGTC Bioproducts (Hessle, UK). Oxyma Pure was obtained from CEM (Buckingham, UK). Dimethylfromimide (DMF), dry DMF, dichloromethane (DCM), diethyl ether and acetonitrile (MeCN) were obtained from Sigma Aldrich. *N*,*N*-Diisopropylcarbodiimide (DIC), trifluoroacetic acid (TFA), HATU and HBTU were obtained from Fluorochem (Hadfield, UK). Triethylamine (TEA), *N*,*N* – diisopropylethylamine (DIPEA), triisopropylsilane (TIPS), piperidine, chloroacetic anhydride, HEPES, Tween 20, sodium chloride (NaCl), magnesium chloride (MgCl2) and sodium hydroxide (NaOH) were obtained from Sigma Aldrich. 1,8-Diazabicyclo[5,4,0]undec-7-ene (DBU) was obtained from Merk (Darmstadt, Germany). Ultra-pure water was produced on-site using a Millipore Elix Q-Gard purification system.

TentaGel S RAM rink amide resin was obtained from Sigma Aldrich. Linear peptide synthesis was performed on Liberty BlueTM HT12 Automated Microwave-assisted Peptide Synthesiser. Samples were sonicated on VWR Ultrasonic Cleaner USC-TH. Sample concentration was performed on Genevac EZ-2 Plus Centrifugal Evaporator. Centrifugation was performed on ISA Ventilated Centrifuge and Eppendorf Centrifuge 5424. Freeze-drying was performed on Alpha 2-4 LD Plus freeze-dryer. Peptide analysis was performed on Waters liquid chromatography – mass spectrometer (LC-MS) system. Peptide purification was performed on Shimadzu LC20-AR high-performance liquid chromatography (HPLC). Surface plasmon resonance assays (SPR) were performed on Biacore S200 (GE Healthcare). Peptide and protein concentrations were determined using the method stated for each experiment. Data were analysed using GraphPad Prism and the instrument-specific software described below.

#### ▪ mRNA library synthesis

$NNK_{6-12}$ DNA libraries used for RaPID selections against RAB27A2S were prepared according to previously published procedures.[1] The library comprised variable regions containing 6–12 consecutive degenerate NNK codons. Individual libraries were combined at a ratio of 0.001:0.032:1:32:33:33:11 for $NNK_6$, $NNK_7$, $NNK_8$, $NNK_9$, $NNK_{10}$, $NNK_{11}$ and $NNK_{12}$, respectively.

DNA libraries were transcribed into mRNA using T7 RNA polymerase for 16h at 37 °C. The resulting transcripts were ligated to a puromycin-containing linker using T4 RNA ligase at 25 °C for 30 min, following previously reported procedures.[1]

#### ▪ Aminoacylation of tRNAs

Flexizymes (eFx and dFx), initiator tRNA fMet_CAU and elongator tRNA Asn_CAU were synthesised according to previously published procedures.[1]

Aminoacylation of the initiator tRNA with *N*-chloroacetyl-D-tyrosine cyanomethyl ester (ClAc-D-Tyr-CME) was performed using 5 mM ClAc-D-Tyr-CME, 600 mM $MgCl_2$, 20% DMSO, 25 μM eFx, 25 μM initiator tRNA and 50 mM HEPES-KOH (pH 7.5). The reaction mixture was incubated on ice for 2h. The aminoacylated tRNA was recovered by ethanol precipitation, and the resulting pellet was washed twice with 70% ethanol containing 0.1 M sodium acetate (pH 5.2), frozen in liquid nitrogen and stored at −80 °C until use.

#### ▪ Preparation of RAB27A2S-immobilised magnetic beads

*N*-terminally biotinylated RAB27A2S was incubated with streptavidin-coated magnetic beads (Invitrogen) for 15 min at 4 °C, immediately before selection, to achieve a protein loading of 0.9 pmol/μL beads.

Remaining unoccupied streptavidin-binding sites were blocked by addition of free biotin to a final concentration of 25 μM, followed by incubation for 15 min at 4 °C. The beads were subsequently washed three times with one bead volume of selection buffer and maintained on ice until use.

Negative-selection beads were prepared using the same procedure, omitting RAB27A2S and incubating the beads with selection buffer alone or supplemented with 25 μM biotin.

#### ▪ RaPID selection against RAB27A2S

RaPID selections against biotinylated RAB27A2S were performed using previously reported procedures.[1] Five consecutive rounds of selection were completed.

mRNA libraries were translated using a custom methionine-deficient Flexible *In vitro* Translation (FIT) system supplemented with ClAc-D-Tyr-tRNA fMet_CAU at a final concentration of 25 μM. The first selection round was performed in a reaction volume of 150 μL, whereas subsequent rounds used 5 μL reactions. Translation reactions were incubated for 30 min at 37 °C, followed by 12 min at 25 °C. Ribosomal complexes were disrupted by addition of EDTA to a final concentration of 18 mM (pH 8.0).

The mRNA–peptide conjugates were reverse transcribed using MMLV reverse transcriptase (RNase H Minus, Promega) at 42 °C for 1 h.

Following reverse transcription, reaction mixtures were buffer exchanged into selection buffer using homemade Sephadex columns (Cytiva). The resulting libraries were supplemented with sheared salmon sperm DNA (1 mg/mL) and acetylated BSA (0.1%) to minimise nonspecific binding.

Three consecutive negative-selection steps were performed using streptavidin beads without immobilised RAB27A2S, each for 30 min at 4 °C. The depleted libraries were subsequently incubated with immobilised biotinylated RAB27A2S for 30 min at 4 °C.

Following selection, beads were washed three times with one bead volume of selection buffer at 4 °C. Bound mRNA/DNA–peptide complexes were recovered by heating the beads in PCR buffer at 95 °C for 5 min.

Recovery of the selected library was quantified by real-time quantitative PCR (qPCR) relative to library standards, negative-selection controls and the input library. Recovered DNA was amplified and used to generate the input library for the subsequent selection round.

After five rounds of selection, double-indexed sequencing libraries were prepared using Nextera XT indices and analysed by next-generation sequencing on an Illumina MiSeq platform using 151-cycle single-end reads.

Sequencing results were ranked according to read abundance and translated into their corresponding peptide sequences for identification of potential RAB27A2S-binding macrocycles.

#### ▪ Peptide Synthesis

Linear peptides were synthesised on Liberty BlueTM HT12 Automated Microwave-assisted Peptide Synthesiser using a standard Fmoc/tBu protocol. The synthesis was conducted on 0.20 mmol scale using TentaGel S RAM rink amide resins (208 mg, 0.24 mmol/g). The resin was swelled in 10 ml of DMF:DCM (1:1, v/v) for 15 min prior to synthesis. Fmoc-protected amino acids were dissolved in DMF to obtain 0.20 M solutions. Fmoc-deprotection solution was prepared as a 20% (v/v) Piperidine 0.1M Oxyma pure solution in DMF. Activator solutions were 0.5M DIC and 0.5M Oxyma Pure in DMF. Washings were performed with DMF and DMF:DCM (1:1, v/v) mixture was used as a transfer solvent. Amino acid coupling and deprotection was carried out according to Liberty BlueTM recommended operation method. At the end of the synthesis, chloroacetylation method was performed using 0.5M solution of chloroacetic anhydride in DMF. The resulting peptides attached to resins were transferred to a fritted syringe and washed with DMF (4ml x3), DCM (4ml x3), dried and stored at -20 °C.

##### Peptide cyclization on resin

The peptide-carrying resins were swelled in 5 ml of DCM for 10 mins. Cys-mmt residues were selectively deprotected using TFA:TIPS:DCM (2.5:5:92.5, v/v/v) washes (4ml x20). The extent of deprotection was judged by the colour-change from orange to colourless. The resin acidity was neutralised with 1% (v/v) TEA in DCM washes (4ml x15) until no fumes were observed. DMF washes (4ml x5) were performed to remove DCM, followed by the treatment with 4ml of DBU:DMF (0.002:3, v/v) for peptide cyclisation. The reaction mixture was left to shake at room temperature for 1.5 hours. The resulting resins carrying cyclised peptides were washed with DMF (4ml x 3), DCM (4ml x 5) and submitted for cleavage.

##### Peptide cleavage

The peptides were fully deprotected and cleaved by treatment with TFA:TIPS:$H_2O$ (95:2.5:2.5, v/v/v) and shaking at room temperature for 1.5 hours. The resins were filtered with cleavage solution washings. The resulting solution was collected and dried under nitrogen flow. The peptides were recovered by precipitation with ice cold diethyl ether. The precipitated peptides were centrifuged at 4000rpm for 2 mins, and the supernatant was discarded. The procedure was repeated four times. The crude linear peptides were dissolved in 5ml of water and submitted for LC-MS analysis. The samples were freeze-dried overnight to obtain porous white solids, which were stored at -20 °C.

##### Peptide cyclisation in solution

After peptide cleavage from the resin, the *N*-chloro-acetylated linear peptides were dissolved in 8 ml of distilled water. The pH was increased to 8-9 using aqueous NaOH (0.1M) solution. The resulting solution was shaken at room temperature for 15 mins. The reaction was quenched by lowering the pH to 3-4 using TFA. The samples were submitted for LC-MS analysis. The cyclised peptides were freeze-dried overnight to obtain porous white solids, which were stored at -20 °C.

##### Peptide LC/MS analysis

The crude peptide samples were analysed on the Waters LC-MS system (2767 sample manager, 2998 UV detector and 3100 mass detector). The mobile phase consisted of 0.1% (v/v) formic acid in water (eluent A) and 0.1% (v/v) formic acid in MeCN (eluent B). The injection volume was 20uL. The eluent gradient started at 5% B and was increased to 98% B over 10 mins. This was followed by 98% B wash for 3 mins, and a sharp decrease to 5% B for equilibration, which was held for further 5 mins. The separation was achieved on a XBridge C18 (100x4.6mm, 5µm). The column temperature was 40 °C and the flow rate was 1.2 mL/min. The ionisation source was electrospray ionisation (ESI) in positive and negative modes. The PDA detector was set for 220-400 nm wavelength range. The expected and observed LC/MS values can be found in the Supplementary Figures.

##### Peptide purification

The dry samples were dissolved in MeCN:Water (5:95, v/v), sonicated for 10min at room temperature and centrifuged for 2 mins at 14680 rpm. The supernatant was collected and filtered through a syringe filter to give a clear solution. The crude peptide samples were purified on the reversed-phase Shimadzu LC20-AR HPLC. The separation was achieved on the Phenomenex AerisTM peptide XB-C18 column (150x21.2mm, 5µm). The mobile phase consisted of 0.1% (v/v) TFA in water (eluent A) and 0.1% (v/v) TFA in MeCN (eluent B). The eluent gradient started at 5% B, held for 2 min, and increased to 40% B over 16 min. This was followed by a sharp increase to 98% B over 2 min, which was held for 2 mins. The gradient was reduced to 5%B over 1 min, which was held until 25 min for column re-equilibration. The column was kept at room temperature, and the flow rate was 20 mL/min. The photodiode array detector was set to 220 nm and 280 nm wavelength. The collected fraction's purity was confirmed using LC-MS. The fractions were concentrated using Genevac EZ-2 Plus Centrifugal Evaporator, then freeze-dried overnight to obtain porous white solids, which were stored at -20 °C.

#### ▪ Protein constructs

The **RAB27A2S** construct comprised human RAB27A (UniProt P51159, residues 1-192) carrying Q78L, C123S and C188S substitutions. The coding sequence was cloned into pET15b vector (Invitrogen) with an N-terminal $His_6$-tag followed by a Tobacco Etch Virus (TEV) protease recognition site (ENLYFQ¦G).

The truncated human **SLP2A** construct (UniProt Q9HCH5) contains the sequence for human Synaptotagmin-like protein 2 (residues 5-62) and was cloned into pET15b vector (Invitrogen) with an N-terminal $His_6$-tag, a GB1 solubility tag sequence (derived from the B1 domain of streptococcal Protein G) and a TEV protease recognition site (ENLYFQ¦G):

HHHHHHSSGLVPRGSHMQYKLILNGKTLKGETTTEAVDAATAEKVFKQYANDNGVDGEWTYDDATKTYTVTE
GGSENLYFQ‖GGTSFLTEEEQEAIMKVLQRDAALKRAEEERVRHLPEKIKDDQQLKNMSGQWFYEAKAKRH.

#### ▪ Protein expression and purification

**RAB27A2S** and **SLP2A** constructs were purified as previously reported.[2,3] Briefly: for protein expression, plasmids were transformed into Escherichia coli BL21 cells and plated on lysogeny-broth (LB) agar containing ampicillin (100 mg/L) for selection. Single colonies were picked for amplification and incubated overnight into LB media containing 100 mg/L Ampicillin at 37 °C, shaking. Large-scale cultures were inoculated at 1-10% v/v and grown at 37 °C until $OD_{600}$ reached approximately 0.7. Protein expression was induced with 0.5 mM isopropyl β-D-1-thiogalactopyranoside (IPTG) for 3 h at 37 °C.

Subsequently cells were harvested at 5,000 × g for 10 min at 4 °C and re-suspended in lysis buffer containing 50 mM Tris at pH 8, 500 mM NaCl and 10 mM imidazole. RAB27A2S buffers additionally contained 5 mM $MgCl_2$. Cells were disrupted at 25kpsi and lysates were clarified at 15,000 × g for 45 min at 4 °C. The supernatant was loaded on a 5 mL HisTrap™ FF Crude $Ni^{2+}$-NTA column equilibrated in lysis buffer. After washing, protein was eluted with 50 mM Tris, pH 8.0, 500 mM NaCl and 300 mM imidazole, with 5 mM $MgCl_2$ for RAB27A2S.

The eluate was buffer exchanged using 2x Cytiva HiTrap™ desalting columns in tandem with desalting buffer 100 mM NaCl, 5 mM $MgCl_2$ and 50 mM Tris, pH 8. Afterwards TEV protease (obtained in-house as previously described[3]) was added to the protein solution at a molar ratio of 1:20 in the presence of 1 mM DTT, and the solution was incubated overnight at 4 °C, shaking. The solution was desalted again with 100 mM NaCl, 5 mM $MgCl_2$ and 50 mM Tris, pH 8 and then loaded on $Ni^{2+}$-NTA resin. The flowthrough was collected, concentrated to 5.5 mg/mL in 150 mM NaCl, 5 mM $MgCl_2$, 20 mM Tris pH 8 buffer. Then, for Rab27A2S only, a 10X buffer containing 10 mM $ZnCl_2$ and 2 M $(NH_4)_2SO_4$, 4 molar excess of GppNHp and 25 units of Antarctic phosphatase (New England Biolabs) were added to the solution and incubated overnight at 4 °C. Finally, the samples were loaded on a superdex S-75 gel filtration column at a flow rate of 1 mL/min. The column was pre-equilibrated with 150 mM NaCl, 5 mM $MgCl_2$, and 20 mM HEPES or TRIS at pH 8 for crystallography. The peaks corresponding to Rab27A2S or SLP2A constructs were assessed by SDS-PAGE, pooled, concentrated and flash-frozen in liquid nitrogen and stored at -80 °C.

#### ▪ Sortase mediated protein labelling

Following TEV cleavage, the *N*-terminal glycine of RAB27A2S or SLP2A was labelled using the evolved Sortase 5M enzyme and a biotin- or TAMRA-containing depsipeptide substrate, respectively, following previously reported methods.[4] RAB27A2S was labelled for immobilisation on streptavidin surfaces (For SPR experiments) and beads (for RaPID selection); SLP2A was labelled to generate the TAMRA-SLP2A tracer used in FP experiments. The protected depsipeptide amino acid Fmoc-Thr(OtBu)-$OCH_2COOH$ was synthesized in two solution-phase steps according to previously published procedure.[4] Semi-preparative LC-MS purification was performed on a Waters system (2767 autosampler, 515 pump, 3100 ESI-MS) and HRMS was performed on a Waters LCT Premier Spectrometer operating in W mode ES +. Rink amide resin (50.0 μmol, 1 eq) was swollen in DMF (2 mL, 30 min), Fmoc deprotected with 20% v/v piperidine in DMF (2 mL, 10 min x 3) and washed sequentially with DMF, DCM and DMF. All coupling steps were carried out with HBTU/DIPEA in 2 mL DMF (30 min x 2). Specifically, FmocThr(OtBu)-$OCH_2COOH$ (3equiv) was activated with HBTU (3equiv) and DIPEA (6equiv); TAMRA (2equiv) with HBTU (2equiv) and DIPEA (4equiv); and E, P, L, A, and biotin (5equiv) with HBTU (5equiv) and DIPEA (10equiv). Crude products were cleaved from the resin with 95% TFA, 2.5% water and 2.5% triisopropylsilane (3 h) and precipitated with cold diethyl ether. The solids were pelleted by centrifugation (15 min, 4300 rpm, 4 °C) and washed three times with diethyl ether. Pelleted products were dried and purified (≥ 95%) by semi-preparative LC-MS equipped with Waters X Select C18 columns running a gradient of MeOH (0.1% FA) in water (0.1% FA) (5-98%, 30 min for the biotin derivative and 20-98%, 30 min for the TAMRA derivative). The products were obtained by lyophilization as bright pink (in case TAMRA derivative) or off-white amorphous solid (in case of biotin derivative). TAMRA-ALPET-Haa was obtained in 46.5% yield (23.2 mg) and Biotin-ALPET-Haa in 32.6% yield (13.3 mg). Purified compounds were characterized by HRMS. The main $[M+H]^+$ peak found for TAMRA-ALPET-Haa was of m/z=813.3799 (calc. 813.3817) and of m/z=999.4451 (calc. 999.4451) for Biotin-ALPET-Haa.

**Labelling of RAB27A2S with Biotin- and/or TAMRA- derivatized peptides:** RAB272S (60 μM) was incubated with the corresponding biotin- or TAMRA-functionalised depsipeptide (180 μM) and Sortase A 5M (9 μM) in reaction buffer containing 50 mM HEPES (pH 7.5), 150 mM NaCl, 5 mM $MgCl_2$ and 5 mM $CaCl_2$. The reaction was carried out at 0 °C for 2 h.

Following incubation, the reaction mixture was subjected to gravity-flow nickel-affinity chromatography using Ni-agarose resin to remove His-tagged Sortase A 5M. The labelled RAB27A2S was collected in the flow-through, and protein purity was assessed by 16% SDS-PAGE.

#### ▪ Surface plasmon resonance (SPR) assay

Surface plasmon resonance (SPR) experiments were performed using a Biacore S200 instrument (GE Healthcare) equipped with a Series S Streptavidin sensor chip. *N*-terminally biotinylated RAB27A2S (200 nM) was immobilised on the streptavidin-coated surface to achieve approximately 400 resonance units (RU). A reference flow cell and buffer-only injections were used for double referencing. Experiments were performed at 25 °C using a running buffer containing 20 mM HEPES (pH 7.4), 150 mM NaCl, 5 mM $MgCl_2$, 0.005% Tween-20 and 0.1% BSA, at a flow rate of 30 μL/min.

Binding affinities of IMP2660 and recombinant SLP2A were determined by injecting increasing concentrations of each ligand (0, 3.9, 7.8, 15.6, 31.25, 62.5, 125, 250, 500, 1000, 2000 and 4000 nM) over immobilised RAB27A2S. SPR responses were recorded in resonance units (RU), and sensorgrams were analysed using the Biacore S200 Evaluation Software. Binding data were fitted using the instrument's affinity analysis model to determine equilibrium dissociation constants ($K_D$). IMP2660 and SLP2A exhibited $K_D$ values of 96 nM and 93 nM, respectively.

#### ▪ Validation of RaPID hits and alanine scan

The macrocyclic peptides identified through RaPID selection (PH1–PH6) were synthesised, purified and evaluated for binding to immobilised biotinylated RAB27A2S by SPR, using the experimental conditions described above. Initial binding experiments were performed using increasing peptide concentrations (0, 62.5, 125, 250, 500, 1000 and 4000 nM). Peptide 1 (PH6) was subsequently evaluated under optimised conditions using single-cycle kinetics with concentrations of 0, 3.9, 7.8, 15.6, 31.25, 62.5, 125, 250, 500 and 1000 nM.

To investigate the contribution of individual amino acid residues to RAB27A2S binding, a series of single-alanine variants of peptide 1 was synthesised and evaluated by SPR under the same experimental conditions. Binding sensorgrams were analysed using the Biacore S200 Evaluation Software, and equilibrium dissociation constants ($K_D$) were determined by fitting the experimental data to a 1:1 binding model. Binding affinities were determined from three independent measurements, and the corresponding $K_D$ values are reported in Supplementary Table S3.

#### SPR Effector binding and competition

Competition between IMP2660 and SLP2A for binding to RAB27A2S was investigated using a Biacore S200 instrument (GE Healthcare). *N*-terminally biotinylated RAB27A2S was immobilised on a Series S Streptavidin sensor chip to approximately 400 resonance units (RU). Experiments were performed at 25 °C in running buffer containing 20 mM HEPES (pH 7.4), 150 mM NaCl, 5 mM $MgCl_2$, 0.005% Tween-20 and 0.1% BSA, at a flow rate of 30 µL/min.

Individual binding experiments were initially performed by injecting serial dilutions of SLP2A or IMP2660 over immobilised RAB27A2S in the absence of competing ligand, yielding dissociation constants ($K_D$) of 93 nM and 96 nM, respectively.

Reciprocal competition experiments were subsequently performed using an A–B–A injection format. In the first experiment, IMP2660 was injected at a fixed concentration of 5 µM, followed by increasing concentrations of SLP2A (0, 3.9, 7.8, 15.6, 31.25, 62.5, 125, 250, 500, 1000, 2000 and 4000 nM) (Figure S5a). Reciprocal experiments were performed using fixed concentrations of SLP2A (5 or 50 µM), with IMP2660 tested over the same concentration range (Figures S5b and S5c).

SPR responses were recorded in resonance units (RU), and competition-associated responses were extracted from the sensorgrams using the Biacore evaluation software. In the presence of IMP2660, SLP2A displayed an apparent $K_D$ of 4.6 µM, compared with 93 nM in the absence of the macrocyclic peptide, consistent with competitive interference with SLP2A binding.

#### Differential scanning fluorimetry

Differential scanning fluorimetry (DSF) was performed to assess the thermal stabilisation of RAB27A2S upon binding to IMP2660. Experiments were conducted using 2.5 µM RAB27A2S in assay buffer containing 20 mM HEPES (pH 7.4), 100 mM NaCl and 5 mM $MgCl_2$, supplemented with 5× SYPRO Orange fluorescent dye (diluted from a 5000× stock). RAB27A2S was analysed in the absence or presence of IMP2660 at final concentrations of 62.5 and 125 µM.

Protein thermal unfolding was monitored by measuring changes in SYPRO Orange fluorescence as a function of temperature. Measurements were performed in triplicate, and fluorescence data were processed to determine the apparent fraction of folded protein. Apparent melting temperatures ($T_m$) were determined from the resulting unfolding curves.

RAB27A2S exhibited an apparent $T_m$ of 53.41 ± 0.05 °C in the absence of IMP2660, which increased to 58.32 ± 0.12 °C and 59.81 ± 0.18 °C in the presence of 62.5 and 125 µM IMP2660, respectively. These corresponded to thermal shifts ($\Delta T_m$) of +4.91 °C and +6.40 °C (Figure 4a and Supplementary Table S4).

#### Fluorescence polarization assay

Fluorescence polarisation (FP) was used to evaluate the ability of IMP2660 to disrupt the interaction between RAB27A2S and its effector protein SLP2A in solution. TAMRA-labelled SLP2A (TAMRA-SLP2A) was used as a fluorescent tracer.

The assay buffer (20 mM HEPES, 100 mM NaCl, 5 mM $MgCl_2$, pH 7.4) was freshly prepared and filtered through a 0.2 µm membrane, followed by the addition of 0.005% Tween-20 and 0.1% BSA.

Binding experiments were performed in black, low-binding, clear-bottom 384-well microplates (Greiner Bio-One, 788096; μCLEAR), with a final reaction volume of 10 μL per well. TAMRA-SLP2A was prepared at a 2× concentration of 20 nM to achieve a final assay concentration of 10 nM.

RAB27A2S was serially diluted two-fold across 24 concentrations, ranging from 5 μM to approximately 0.6 pM and combined with TAMRA-SLP2A in the absence or presence of fixed final concentrations of IMP2660 (100 nM, 1 μM or 10 μM). The assay was performed in triplicate, with a final reaction volume of 10 μL per well.

Fluorescence polarisation was measured using a CLARIOstar plate reader equipped with a TAMRA FP filter set (excitation 540 ± 10 nm, emission 590 ± 10 nm; 20 flashes per well). Data were analysed using GraphPad Prism (version 10.2.3). Binding curves were fitted by nonlinear regression using a four-parameter logistic model (log[concentration] versus response, variable slope), with the upper and lower plateaus shared across datasets. Apparent dissociation constants ($K_D$) were derived from the fitted curves. In the absence of IMP2660, TAMRA-SLP2A bound RAB27A2S with an apparent $K_D$ of 40 nM. Increasing concentrations of IMP2660 resulted in a concentration-dependent rightward shift in the binding curves, yielding apparent $K_D$ values of 55 nM, 275 nM and 1.8 μM at 100 nM, 1 μM and 10 μM IMP2660, respectively (Table S5).

- ## Protein crystallisation

Pure samples of Rab27A2S (10 mg/mL) were incubated overnight at 4 °C with 30 μM IMP2660 then cleared by centrifugation at 13k RPM. Following a small screening in 96-well plate, a single crystal was obtained at 4 °C using the sitting-drop vapor-diffusion method (200 nL protein + 200 nL mother liquor) with a mother liquor containing 0.15 M Ammonium sulfate, 0.1 M MES pH 6, 15% w/v PEG 4000.

- ## Crystal diffraction, data collection and data processing

Data collections were carried out at Diamond Light Source in Oxford using the i24 beamline at 100K, wavelength 0.9999 Å, using Pilatus detector. Data processing was carried out using Dials (ref). Initial phases were calculated using the molecular replacement program phaser. The coordinates of Rab27a from chain A of the Rab27a-Slp2a complex (PDB:3BC1) without the nucleotide and the magnesium ion were used as the search model. Subsequently, the initial model generated by phaser was refined through an iterative cycle using COOT and REFMAC5. Final model structures were validated using the Molprobity server at http://molprobity.biochem.duke.edu. All structure images were prepared using ChimeraX or Pymol. X-ray data collection, processing and refinement statistics are given in Table S1.

### Appendix

<u>LC-MS traces and mass spec of the HPLC-purified cyclic peptides reported in the manuscript</u>

**Compound 1 – V3A**

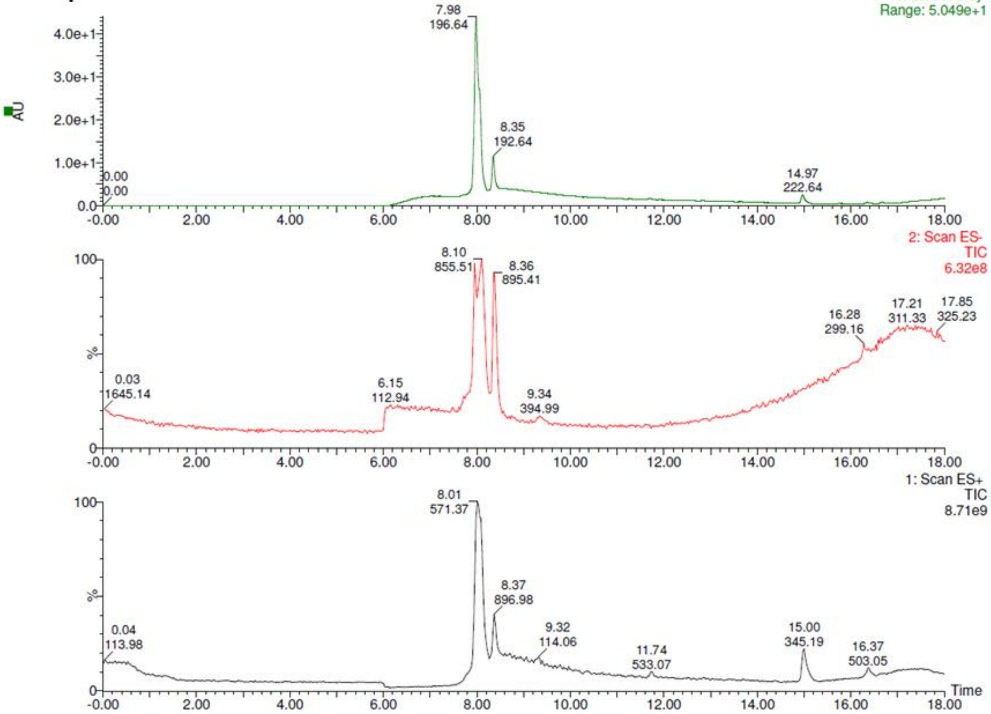


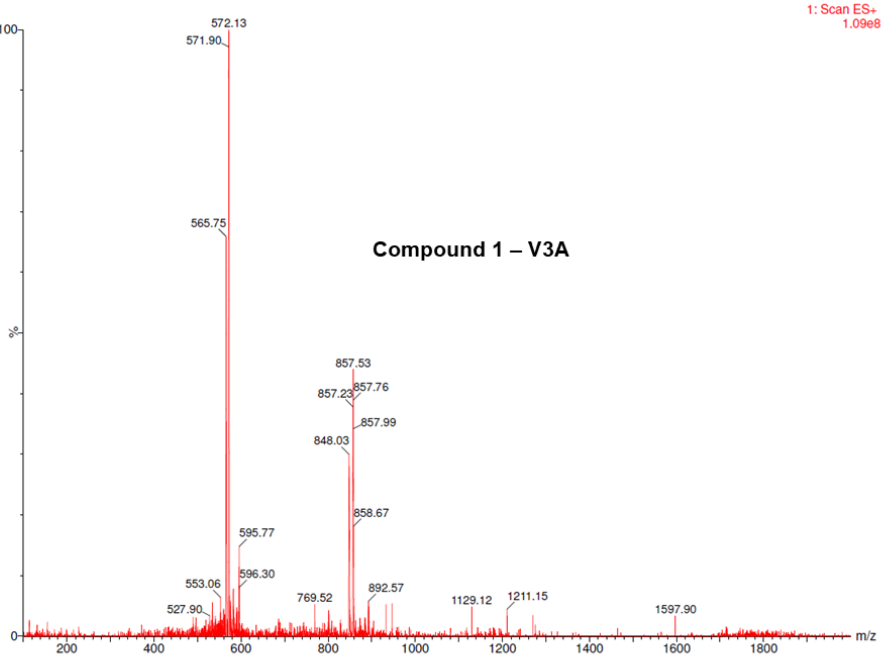

**Compound 1 – G4A**

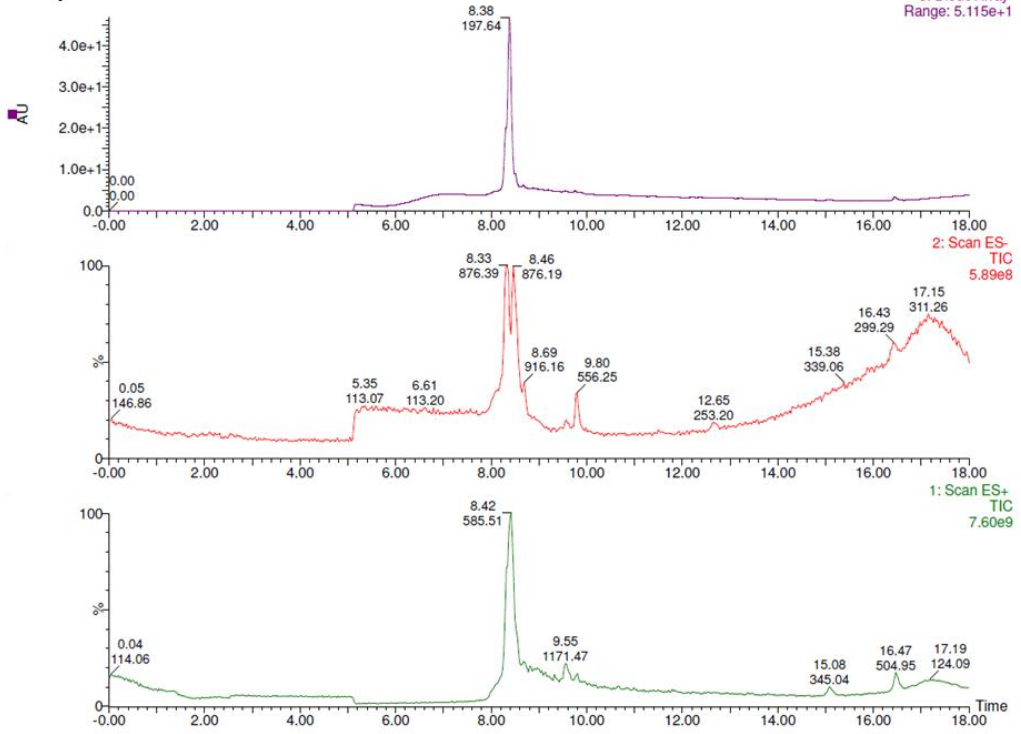


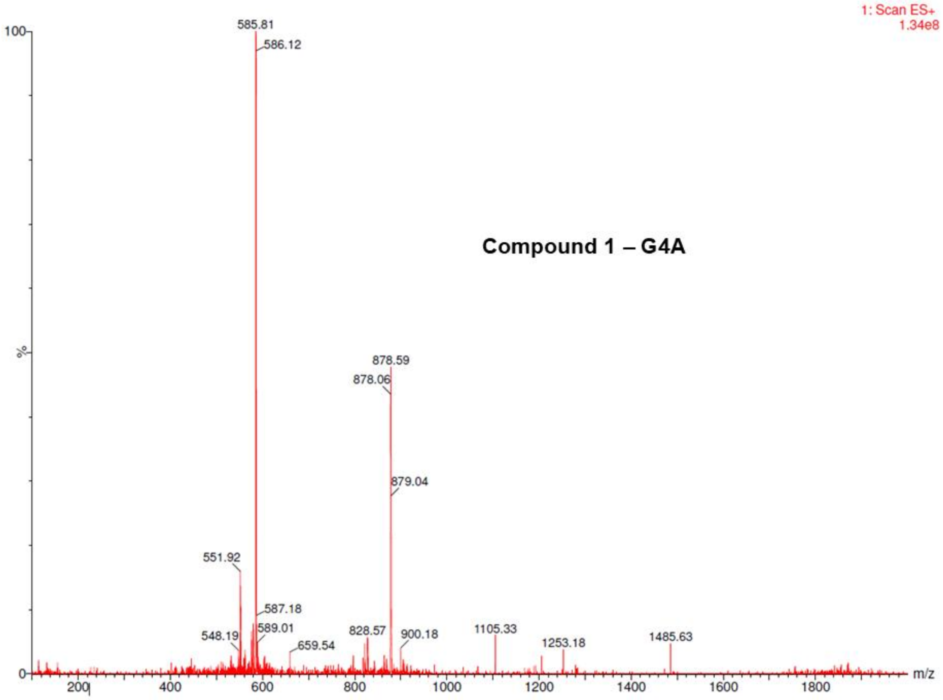

**Compound 1 – S5A**

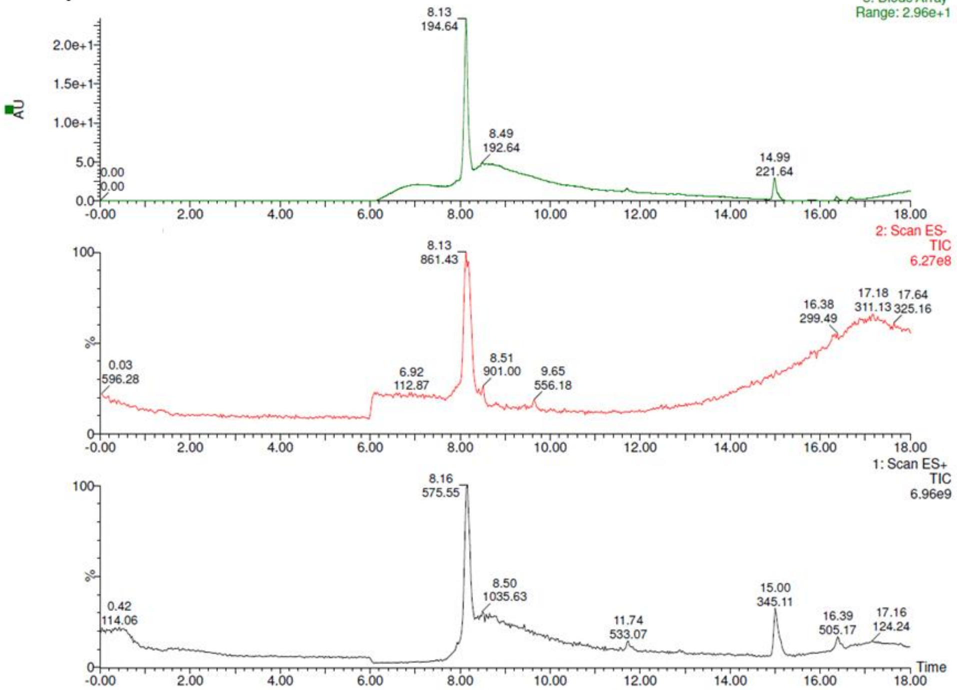


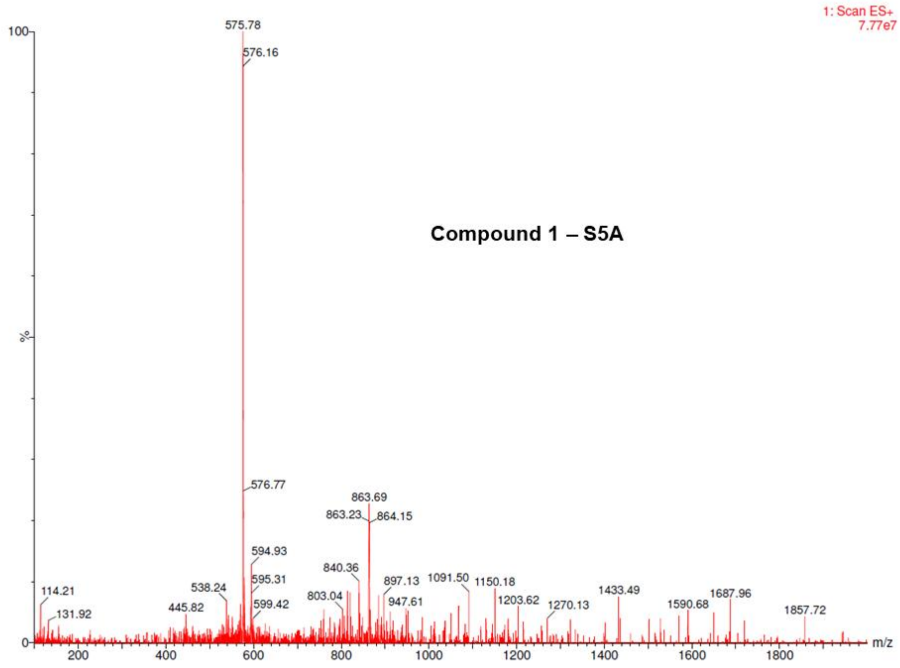

#### Compound 1 – T8A

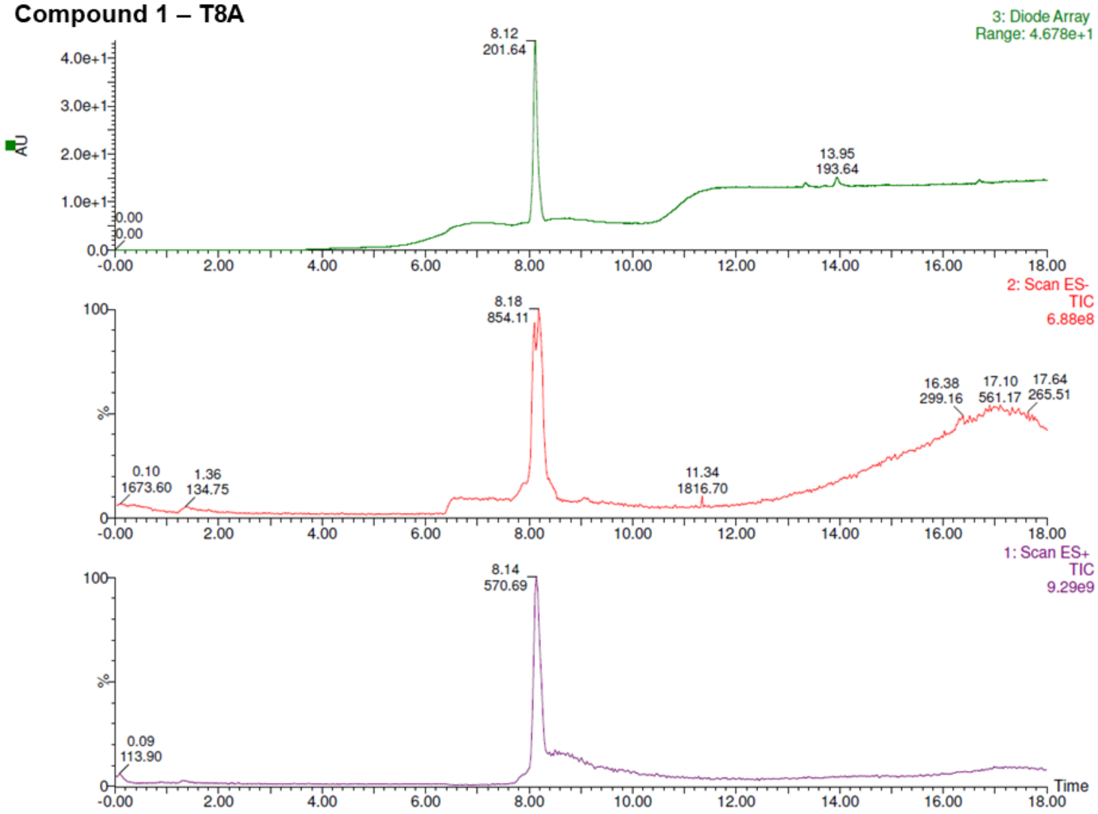


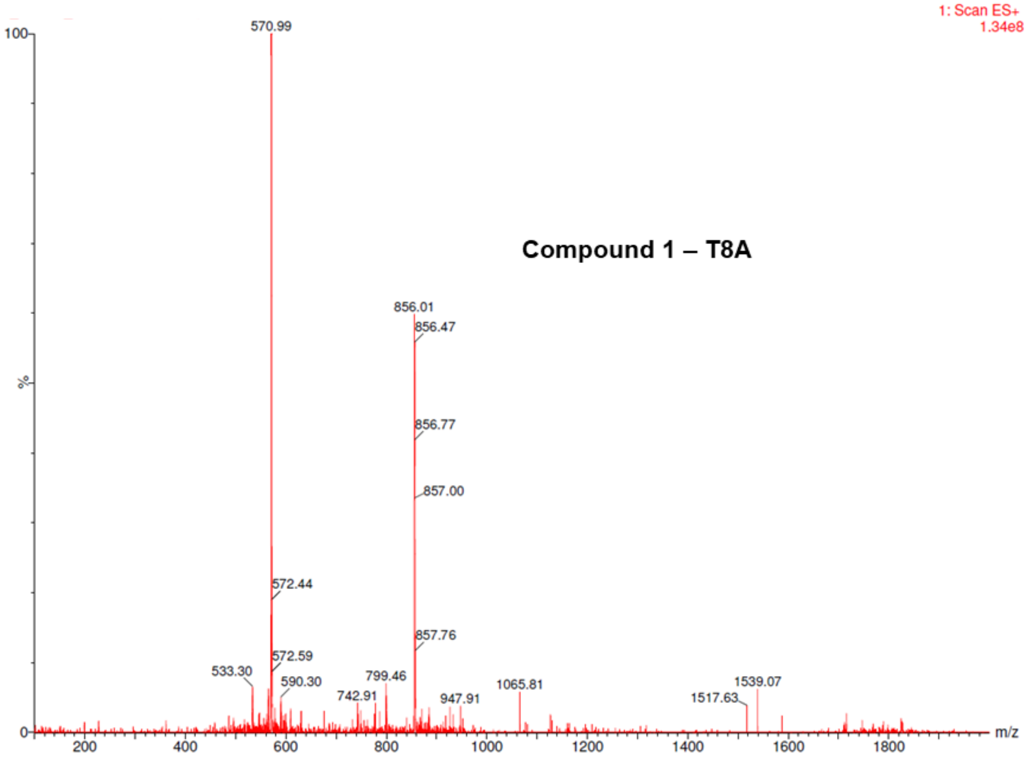

**Compound 1 – I9A**

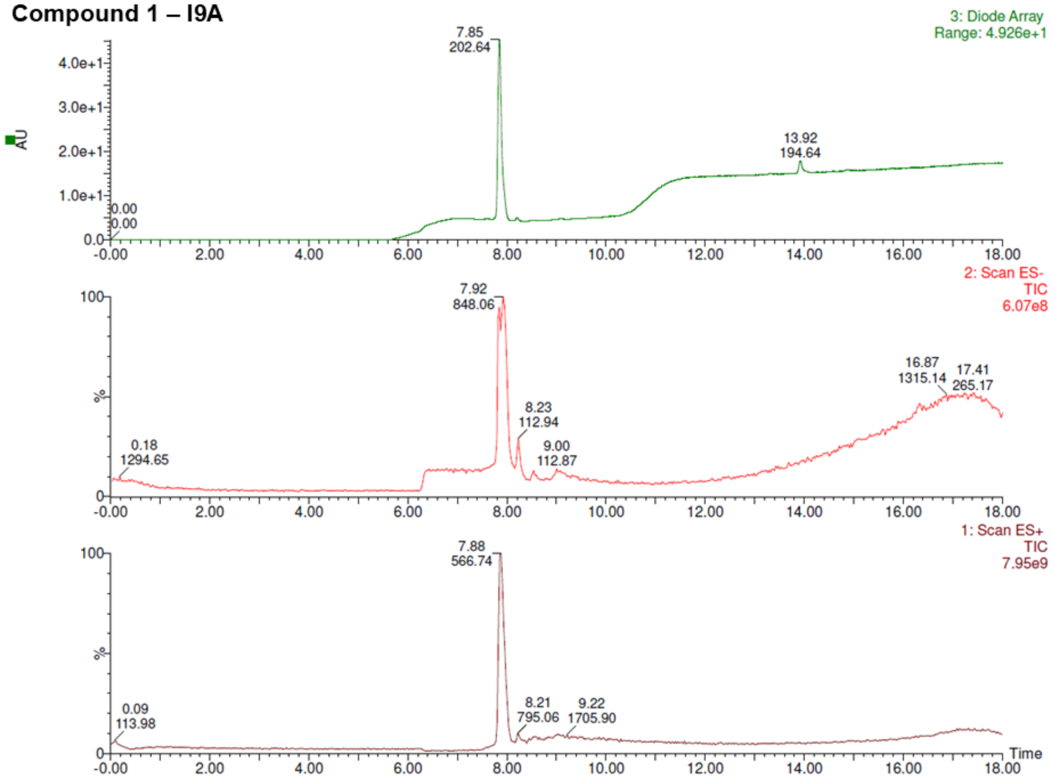


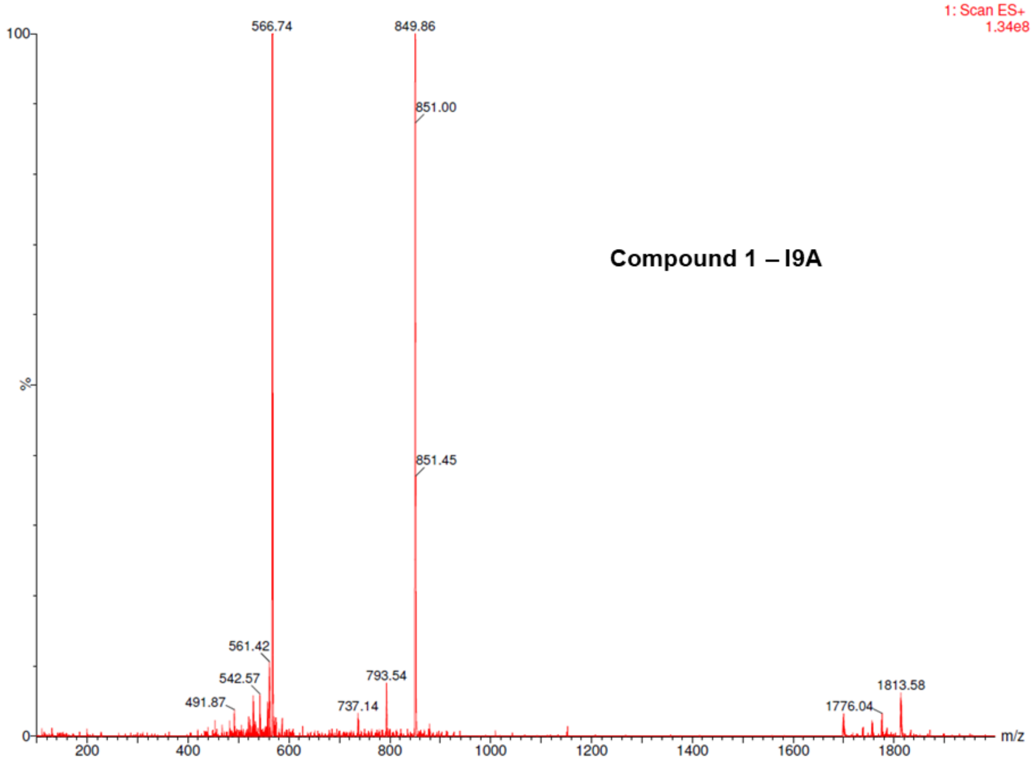
